# Different metabolite profiles across *Penicillium roqueforti* populations associated with ecological niche specialisation and domestication

**DOI:** 10.1101/2024.01.12.575369

**Authors:** Ewen Crequer, Emmanuel Coton, Gwennina Cueff, Johan V. Cristiansen, Jens C. Frisvad, Ricardo Rodriguez de la Vega, Tatiana Giraud, Jean-Luc Jany, Monika Coton

## Abstract

Fungi are known to produce many chemically diversified metabolites, yet their ecological roles are not always fully understood. The blue cheese making fungus *Penicillium roqueforti* thrives in different ecological niches and is known to produce a wide range of metabolites, including mycotoxins. Three *P. roqueforti* populations have been domesticated for cheese production and two populations thrive in other anthropized environments, i.e., spoiled food, lumber and silage. Here, we looked for differences in targeted and untargeted metabolite production profiles between populations using HPLC-HR-Q-TOF and UHPLC-Q-TOF-HR-MS/MS. The non-cheese populations produced several fatty acids and different terpenoids, lacking in cheese strains. The Termignon cheese population displayed intermediate metabolite profiles between cheese and non-cheese populations, as previously shown for other traits. The non-Roquefort population, the cheese population with the strongest domestication syndrome, produced the lowest quantities of measured metabolites, including known mycotoxins such as mycophenolic acid (MPA), andrastin A and PR toxin. Its inability to produce MPA was due to a deletion in the *mpaC* gene, while a premature stop codon in ORF 11 of the PR toxin gene cluster explained its absence and the accumulation of its eremofortin A & B intermediates. In the Roquefort population, we detected no PR toxin nor eremofortins A or B, but found no indel or frameshift mutation, suggesting downregulation. Our results suggest that domesticated cheese populations were selected for lower toxin production while populations from other anthropized environments maintained high metabolite diversity, the bioactivities of these compounds being likely important in these ecological niches.

## Introduction

Fungi are known to produce a wide range of chemically diversified metabolites in nature, that are key for fungal development, interactions, survival and/or warfare with other microorganisms in complex ecosystems. These metabolites include toxins, antimicrobial compounds, bioactive compounds, e.g antibiotics, anti-cancer and immunosuppressants, as well as molecules involved in communication or protection from UV damage (Keller 2019; Stroe et al. 2023). The role of metabolites is however still not fully understood and identifying different metabolite profiles in populations thriving in distinct niches may contribute to our understanding of their ecological role.

*Penicillium roqueforti* is a highly interesting filamentous fungus from an ecological point of view, as it colonises a multitude of niches thus highlighting its ability to adapt to different environments. This species is especially well known worldwide for its beneficial role in blue cheese production (Gillot et al. 2015, 2017b; Dumas et al. 2020), but has also been isolated from lumber and found as a common contaminant in silage or food products like dairy, fruits and bakery (Pitt and Hocking 2009; Crequer et al. 2023). *Penicillium roqueforti* produces many chemically diverse metabolites, including many so called secondary metabolites, now rather referred to as specialised metabolites, with known bioactive properties. Some of these metabolites correspond to mycotoxins, the most toxic being the aristolochene-derived sesquiterpene PR toxin (*P. roqueforti* toxin), which can be a threat to feed and food safety. For example, PR toxin in silage causes liver toxicity or subacute symptoms in livestock (Gallo et al. 2015; Hymery et al. 2017; Dubey et al. 2018). In cheese, PR toxin is considered unstable (Scott and Kennedy 1976) and apparently degraded to PR imine (Siemens and Zawistowski 1993), a molecule with lower toxicity (Hymery et al. 2014). *Penicillium roqueforti* also produces the alkaloid mycotoxin roquefortine C (ROQ C) and meroterpenoid compound mycophenolic acid (MPA), which can be found in various cheeses in a wide range of concentrations (Scott and Kennedy 1976; Lafont et al. 1979; Engel et al. 1982; Finoli et al. 2001; Kokkonen et al. 2005; Usleber et al. 2008; Fontaine et al. 2015). These compounds have relatively low cytotoxic effects compared to other mycotoxins with regulatory limits in foods in the EU (European Union), such as aflatoxins, ochratoxin A or patulin (Commission Regulation EU N° 2023/905) (Fontaine et al. 2016). Mycophenolic acid is even widely used in the medical field as a treatment to prevent organ transplant rejection. Andrastin A (AND A) is another *P. roqueforti* meroterpenoid of interest, with promising anticancer activity (Nielsen et al. 2005). Additional metabolites may also have contributed to *P. roqueforti* adaptation to its environment, *e.g.* clavines, terpenoids, alkaloids or peptides. For example, two tetrapeptides, Phe-Val-Val-Phe and Phe-Val-Val-Tyr, likely have potential antimicrobial properties (Hammerl et al. 2019). Studying the differences in metabolite production between *P. roqueforti* populations may help gain a more general understanding of their ecological role in diverse niches.

Five populations have indeed been identified in *P. roqueforti*, corresponding to three cheese populations and two non-cheese populations (Dumas et al. 2020; Crequer et al. 2023). A first cheese population, called non-Roquefort, corresponds to a clonal lineage (Dumas et al. 2020), and is used worldwide for the production of most kinds of blue cheeses (*e.g.* Gorgonzola, Cabrales, Stilton and Danablue). This population presents numerous beneficial traits for large-scale cheese making, such as higher salt and lactic acid tolerance, faster growth on cheese and faster lipolysis (Dumas et al. 2020; Caron et al. 2021; Crequer et al. 2023). A second cheese population is mainly associated with the Roquefort protected designation of origin (PDO) (Gillot et al. 2015; Dumas et al. 2020). This Roquefort population harbours a bit more genetic diversity and displays traits beneficial for cheese making following more traditional processes, such as longer conservation and culture on bread (Dumas et al. 2020). More recently, a third cheese population was identified, in Termignon blue cheeses that are not inoculated with *P. roqueforti* spores, that instead spontaneously colonise these specific cheeses in the French Alps from the environment. These Termignon strains exhibit intermediary phenotypic traits between the cheese and non-cheese populations and likely correspond to descendants of an ancient domesticated population with mild domestication syndrome (Crequer et al. 2023). Two genetically different non-cheese populations have been identified, one mostly associated with silage, as well as to a lesser extent spoiled food, and the other with spoiled food and lumber (Dumas et al. 2020). These populations have much higher genetic diversity than the cheese populations and have differentiated from each other more recently than from the cheese populations (Dumas et al. 2020).

These genetically differentiated populations of *P. roqueforti* thriving in different environments constitute a great model for studying adaptation to different substrates and in particular the potential ecological roles of the metabolites they produce, in cheese versus other anthropized environments. The non-Roquefort and Roquefort cheese populations result from two different domestication events (Dumas et al. 2020), in the context of more industrial and more traditional production processes, respectively, so that selection on metabolite production or the underlying genetic mechanisms may have been different. Here, we therefore compared the metabolite production profiles between the five *P. roqueforti* populations using targeted (for seven known metabolites, including mycotoxins) and untargeted metabolomics. We also explored the genetic mechanisms underlying the differences using available genomes.

## 2. Material and methods

### Strain collection and conidium suspension preparation

For metabolite profiling, we randomly chose 44 strains from the five known *P. roqueforti* populations (Dumas et al. 2020; Crequer et al. 2023): twelve strains from the non-Roquefort population, eight from the Roquefort population and ten from each of the lumber/spoiled food and the silage/spoiled food populations. We also used the four available strains sampled from Termignon blue cheeses. All strains are available in the ESE (Ecology Systematics and Evolution, Paris Saclay university) or UBOCC (https://nouveau.univ-brest.fr/ubocc/fr) culture collections (Suppl. Table S1).

Conidium suspensions were prepared for the various experiments by cultivating the fungal strains for six days at 25°C on potato dextrose agar (PDA, Difco, Fisher Scientific). Two mL of Tween 80 (0.045 %, v/v) were then added on each plate and conidia were scraped off the surface. Conidium concentrations in the suspensions were estimated using Malassez cells and adjusted to 5.10^5^ conidia.mL^−1^ with Tween 80, in 20% glycerol. Suspensions were then stored at −80°C for cultures for metabolite extraction.

### Metabolite extraction

For metabolite production measurements, we grew fungal cultures in 24-well sterile microplates containing two mL of yeast extract sucrose (YES) agar medium buffered at pH 4.5 with phosphate-citrate buffer and characterised by a high C/N ratio that increases metabolite production in *Penicillium* fungi (Frisvad and Filtenborg 1983). For each strain, 2 µL of the previously prepared spore suspension was inoculated in the centre of the well. Six replicates per strain were performed, three for secondary metabolite analyses and three for fungal dry-weight measurements. In the latter case, growth was performed on cellophane disks to collect fungal mycelium. The plates were incubated at 25°C in the dark for ten days and then stored at −20°C until metabolite profiling. For metabolite extractions, we used an optimised high-throughput extraction method (Gillot et al. 2017b; Lo et al. 2023). After thawing, we homogenised 2 g aliquots (the entire YES culture with mould obtained from a well) with a sterile flat spatula, to which we added 12.5 mL of acetonitrile (ACN) supplemented with 0.1% formic acid (v/v); samples were then vortexed for 30 sec followed by 15 min sonication. Extracts were again vortexed before centrifugation for 10 min at 5000 g at 4°C. The supernatants were directly collected and filtered through 0.45 µm polytetrafluoroethylene membrane filters into amber vials and stored at −20°C until analysis.

#### Targeted secondary metabolite detection and quantification

Targeted secondary metabolite characteristics used for quantifications are given in Table S3 and included commercially available extrolites produced by *Penicillium* species: andrastin A (AND A), eremofortins A & B (ERE A & B), (iso)-fumigaclavine A (FUM A), mycophenolic acid (MPA) and roquefortin C (ROQ C). AND A, ERE A & B and FUM A standards were obtained from Bioviotica (Goettingen, Germany), and others from Sigma-Aldrich (St Louis, MO, USA). All stock solutions were prepared in dimethyl sulfoxide (DMSO) at 1 mg.mL^−1^ in amber vials. For these analyses, metabolite identification was performed using both the mean retention time ± 1 min and the corresponding ions listed in (Supp. Table S2). We used a matrix-matched calibration curve for reliable secondary metabolite quantification with final concentrations ranging from 1 to 10000 ng.mL^−1^ according to the target metabolite and method performance was carried out as previously described (Gillot et al. 2017b). All metabolite concentrations were determined using the Agilent MassHunter Workstation Software (Agilent Technologies, Sanat Clara, CA, USA) with a linear regression model. Specific mycotoxin production was expressed as ng per g of extracted matrix and mg of fungal dry weight (ng.g^−1^.mg^−1^). For PR toxin, a purified solution with unknown concentration was previously obtained (Gillot et al. 2017b) and diluted 1X, 2X, and 5X (taking into account matrix effect) to ensure peak separation and determine the detection limit. According to International Council for Harmonisation guidelines (ICH Harmonized Tripartite Guideline, 2005), detection limit and quantification limit of each metabolite was obtained by multiplying the standard deviation of y intercepts of regression lines divided by the slope, by 3.3 and 10, respectively. Means per strain across replicates (supp. Tables S3 and S4) were used to compare populations. Electrospray ionization (+/-) modes were both systematically performed on all samples. Method performance characteristics were obtained for the seven targeted metabolites prepared in blank extraction of YES medium and all seven metabolites were eluted at different retention times including MPA and PR toxin known to have a quantifying ion with the same mass (m/z 321.13; Supp. Table S2). Linearity (R^2^) for each standard curve was determined to be above 0.982 (Supp. Table S2) for all metabolites in electrospray ionisation positive mode, ESI +, with better performance for detection in ESI + than ESI −.

Targeted analyses were performed on an Agilent 6530 Accurate-Mass Quadrupole Time-of-Flight mass spectrometry system equipped with a binary pump 1260 and degasser (Q-TOF LC/MS), well plate autosampler set to 10°C and a thermostatted column compartment. Filtered 2 µL aliquots were injected into a ZORBAX Extend C-18 column (2.1×50 mm and 1.8 µm, 600 bar) maintained at 35°C with a flow rate set to 0.3 mL.min^−1^. The mobile phase A contained milli-Q water + 0.1% formic acid (v/v) and 0.1% ammonium formate (v/v) while mobile phase B was ACN + 0.1% formic acid. Mobile phase B was maintained at 10% for 4 min followed by a gradient from 10 to 100% for 36 min. Then, mobile phase B was maintained at 100% for 5 min before a 5 min post-time. Samples were ionised in both positive (ESI+) and negative (ESI−) electrospray ionisation modes in the mass spectrometer with the following parameters: capillary voltage 4 kV, source temperature 325°C, nebulizer pressure 50 psig, drying gas 12 L.min^−1^, ion range 100-1000 m/z.

#### Untargeted metabolite analysis, data processing and metabolite identifications

##### UHPLC-Q-TOF-HRMS/MS analysis

We also used an untargeted metabolomics approach with an ultra-high-performance liquid chromatography - diode array detection - quadrupole time of flight mass spectrometry (UHPLC-DAD-Q-TOF-MS/MS) during metabolite profiling on three *P. roqueforti* extracts, the lumber L6, Roquefort R3 and Termignon T4 strains (biological triplicates). These extracts were selected based on the above mentioned targeted LC-QTOF analyses as they also displayed multiple unknown compounds and the three extracts covered the full spectrum of these observed unknown metabolites.

We detected metabolites using an Agilent 6545 Quadrupole Time-of-Flight (Q-TOF) MS equipped with an UHPLC Agilent Infinity 1290 (Agilent Technologies, Santa Clara, CA,502 USA) including a diode array detector. Separation was done on a Poroshell 120 Phenyl Hexyl column (150 × 2.1 mm i.d., 1.9 μm; Agilent Technologies, Santa Clara, CA) maintained at 40°C. Samples injected (1 μL) were eluted with a flow rate set to 0.35 mL min−1 using a linear gradient increasing from 10% acetonitrile (LC-MS grade) in Milli-Q water supplemented with 20 mM formic acid to 100% over the first 10 min, maintaining 100% for 2 min before decreasing back to 10% in 0.1 min and holding initial conditions for 3 min before the next run. The Agilent accurate-mass 6530 Quadrupole Time-of-Flight (Q-TOF) liquid chromatography/mass spectrometer (LC/MS) system was equipped with an Agilent Dual Jet Stream electrospray ion source (ESI) with a drying gas temperature set to 250°C and flow of 12 L.min^−1^. Samples were ionised in positive (ESI+) electrospray ionisation modes in the mass spectrometer with the following parameters: capillary voltage 4 kV, nozzle voltage 500 V, ion range 100-1000 m/z and auto MS/MS fragmentation at three collision energies (10, 20 and 40 eV). The acquisition rate was set to 10 spectra per second and MS spectra were recorded as centroid data. Reference masses (two [M+H]+ ions were: 186.2216 and 922.0098) were injected in the second sprayer using a supplementary LC pump at 15µl.min-1 flow rate using a 1:100 splitter.

##### LC-MS/MS data processing

The generated mass spectrometry (MS) data, recorded as centroid data, were analysed using both an in-house library search with the Agilent MassHunter PCDL manager and MZmine3, GNPS (global natural product social molecular networking) and SIRIUS tools available on GitHub (https://github.com). For MZmine3 analyses, centroid data were converted into the community standard for mass spectrometry data, mzML, using the ProteoWizard software (version 3.0.22112, MSConvert tool) (Martens et al. 2011; Chambers et al. 2012). MZmine3 software (Schmid et al. 2023) was then used to process mzML files and the batch file that was created included the different processing steps as feature detection, deconvolution and filtering. The final feature quantification table, exported in the MGF (mascot generic format) standard format, was used in the GNPS Networking web-based mass spectrometry ecosystem (https://gnps.ucsd.edu/) to generate a molecular network for feature determinations using Cytoscape v3.9.1 (https://cytoscape.org/) and SIRIUS 5.6.3 (https://bio.informatik.uni-jena.de/software/sirius/; (Dührkop et al. 2019)) softwares for further downstream analyses and chemical family predictions and/or identifications.

After data processing of UHPLC-Q-TOF-MS/MS spectra, the LC-Q-TOF/MS spectra for the same three strains were also analysed using SIRIUS 5.6.3 (Dührkop et al. 2019) and the two data sets were compared to match both data sets together (*i.e.* identify common ions), determine putative formulas and compare MS-MS spectra against databases available in SIRIUS to predict the metabolite family and, when possible, identify metabolites. This comparison allowed us to then transpose this data to all the generated metabolite data and extract identified ion peak areas from all LC-Q-TOF spectra. We were therefore able to obtain the metabolite production profile per strain, which was used to compare metabolite production profiles between populations. Control samples were also included, corresponding to the blank YES medium (no mould); it was used to extract and distinguish compounds produced by strains from compounds originally present in the medium.

### Statistical analyses

Statistical analyses for testing differences in metabolite production between populations were performed using the R software (version 4.2.1, https://www.r-project.org/). Shapiro-Wilk and Bartlett tests (package *rstatix*, R) were performed to assess normality and homoscedasticity of residuals in each population. If the data, the racine-transformed data or the log-transformed data did not deviate from normality, populations were compared using ANOVA type I and Tukey tests were used as *post-hoc* tests. If the data, the log-transformed data and square-root transformed data significantly deviated from normality, a Kruskall-Wallis test was performed on raw data to compare populations, followed by Dunn tests as *post-hoc* tests (Suppl. Table. S5).

### Comparative analysis of secondary metabolite biosynthetic gene clusters

In order to study the targeted metabolite biosynthetic gene clusters, we used the annotations of the LCP06136 (Caron, *et al*., 2024) and FM164 reference genomes (genbank accession numbers GCA_000513255.1), and we used the genomes of all other strains analysed here that were previous assembled from Illumina data (Dumas et al. 2020; Crequer et al. 2023). We lifted the annotations of the known gene clusters controlling the production of the studied metabolites, MPA, PR toxin, FUM A, AND A and ROQ C, from the two reference genomes to the Illumina genomes (accession numbers in Table S1), for all the phenotyped strains, using *liftoff* v1.6.3 (Shumate and Salzberg 2021). Protein sequences and coding DNA sequences (CDS) of each gene were extracted from each genome using *gffread* v0.12.1 (Pertea and Pertea 2020). Reference gene annotation used corresponded to the annotation with the longest CDS, to be conservative regarding the likelihood to detect complete genes. Protein and CDS sequences were aligned using *mafft* v7.475 (Katoh and Standley 2013) and analysed using *Jalview* 2.11.2.0 (Waterhouse et al. 2009; Troshin et al. 2018).

## 3. Results

### Metabolite profiles are different between *P. roqueforti* populations

We performed targeted and untargeted LC-Q-TOF metabolite profiling on 44 *P. roqueforti* strains from the five identified *P. roqueforti* populations (Suppl. Table 1), and identified metabolites of interest using in-house local databases or *in silico* Global Natural Products Social Molecular Networking (GNPS) molecular networks and SIRIUS classification. Based on local databases, we identified 16 metabolites (Table 1): agroclavine, AND A, FUM A, MPA-associated molecules (MPA, MPA isomer, homo MPA, MPA prenyl and MPA IV), roquefortines C and D, PR toxin, eremofortins A and B, and tetrapetides, i.e., cyclo(Phe-Val-Val-Phe), Phe-Val-Val-Phe and Phe-Val-Val-Tyr. We also predicted the chemical classes for 20 other metabolites, including eight fatty acids, among which some fatty acid amides (*e.g.* putative lauryldiethanolamine), 11 terpenoids, among which a putative eremofortin C, and one alkaloid (putative festuclavine), while 11 others remained as unknown metabolites (Table 1). While some metabolites were produced by all strains in all five *P. roqueforti* populations, others varied either qualitatively or quantitatively. Principal component analysis (PCA) performed on all 47 metabolites separated the different *P. roqueforti* populations (Figure 1). The two first dimensions explained 45.94% of the variation, while dimensions 3 and 4 explained 17.5% of the variance. The first dimension separated the cheese populations from the non-cheese populations. Dimension 1 was positively associated in particular with PR toxin, AND A, fatty acids and terpenes, and negatively associated with ROQ C and D, and Phe-Val-Val-Phe. The second PCA dimension mainly separated the non-Roquefort population from the two other cheese populations based on positive associations with ERE A & B, fatty amide 2 and terpene 3 and negative ones with MPA and MPA-associated metabolites (MPA isomer, homo MPA, MPA prenyl and MPA IV), as well as the unknown molecules 4, 8, 11, 16, 17 and 19. The third PCA axis separated the Roquefort populations from all other populations (Figure 1) and was positively associated with ROQ C, ROQ D and the unknown metabolites 19, 20 and 21. The fourth dimension separated the silage/spoiled food population from the other ones and was positively associated with several fatty acids (*e.g.* 1, 5, 7 and 9 and putative lauryldiethanolamine).

**Figure 1:**
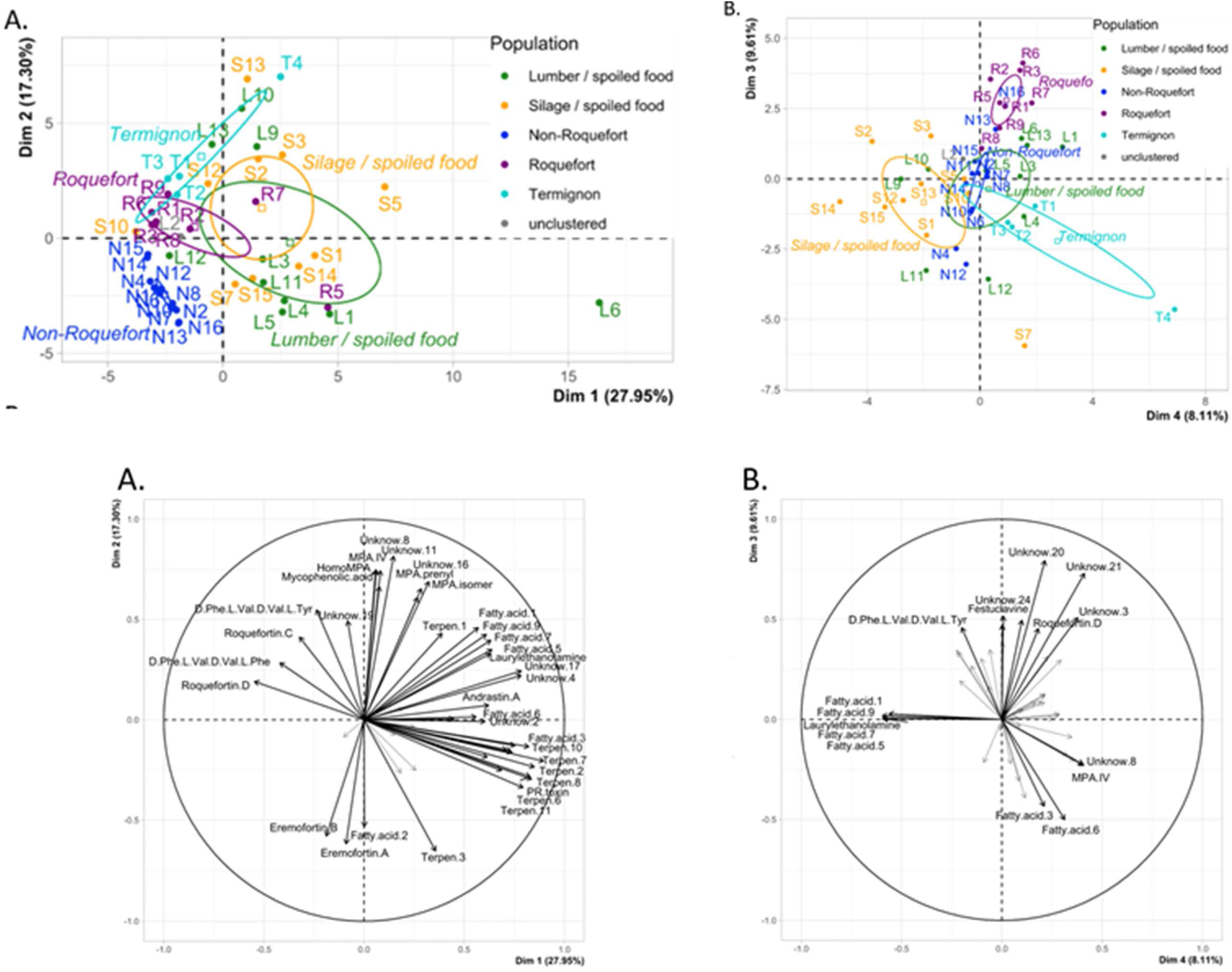
Principal component analysis (PCA) illustrating the metabolite profile differences between *Penicillium roqueforti* populations. **A.** Strains on the first two axes of the PCA. **B**. strains on the third and fourth axes of the PCA. In **A**. and **B**. a confidence ellipse at 95% is drawn for each of the five populations. The percentage of variance explained by the axes are indicated. The same colour code is used as in the other figures: green for the lumber/spoiled food population, orange for the silage/spoiled food population, dark blue for the non-Roquefort cheese population, purple for the Roquefort cheese population and light blue for the Termignon cheese population. The strain IDs are provided in Suppl. Table 1. **C**. Association between the two first and **D.** the third and fourth PCA axes and the variables which corresponded to the selected metabolites from metabolite profiling of 44 *P. roqueforti* strains.

**Table 1.**
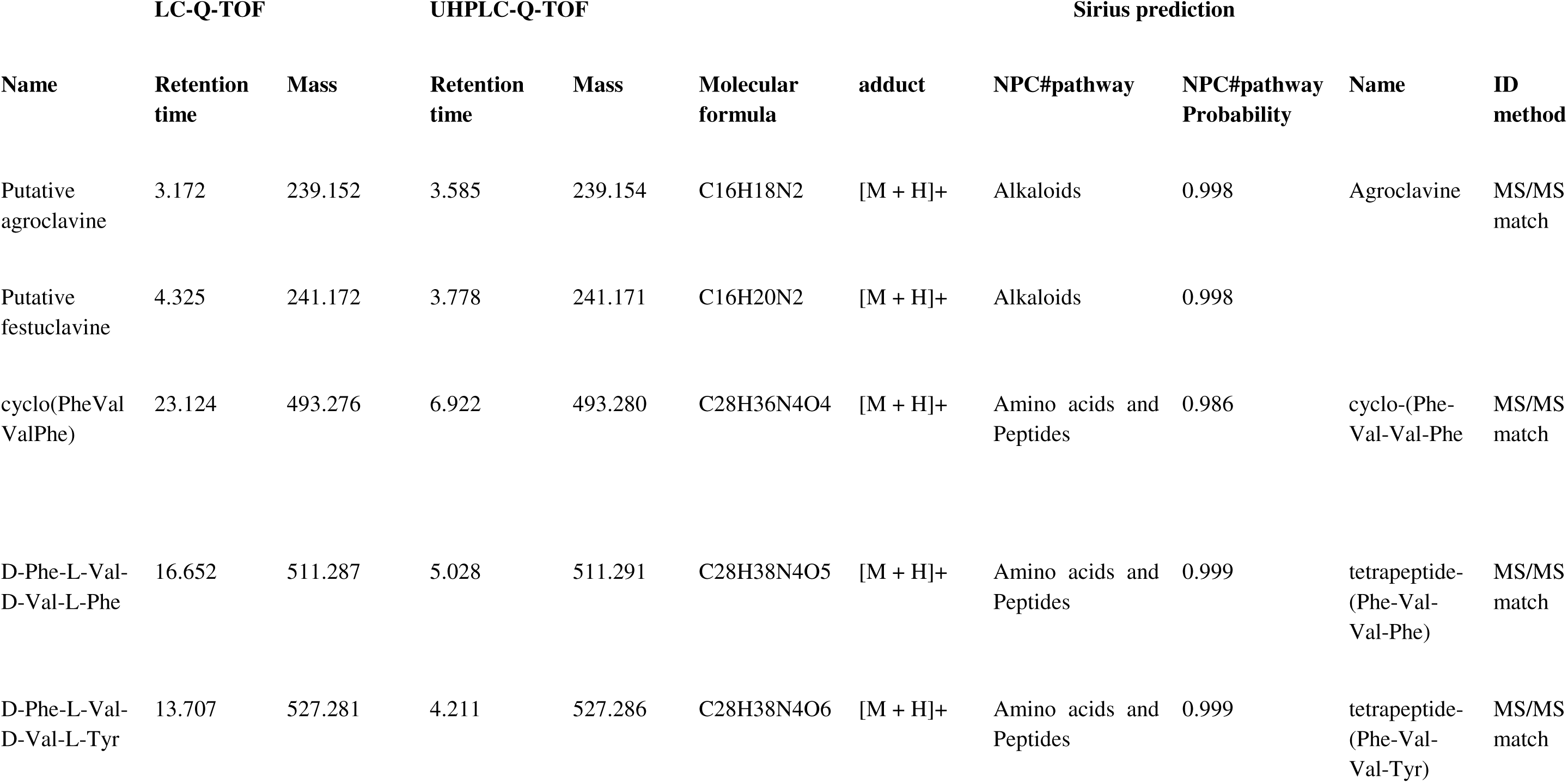
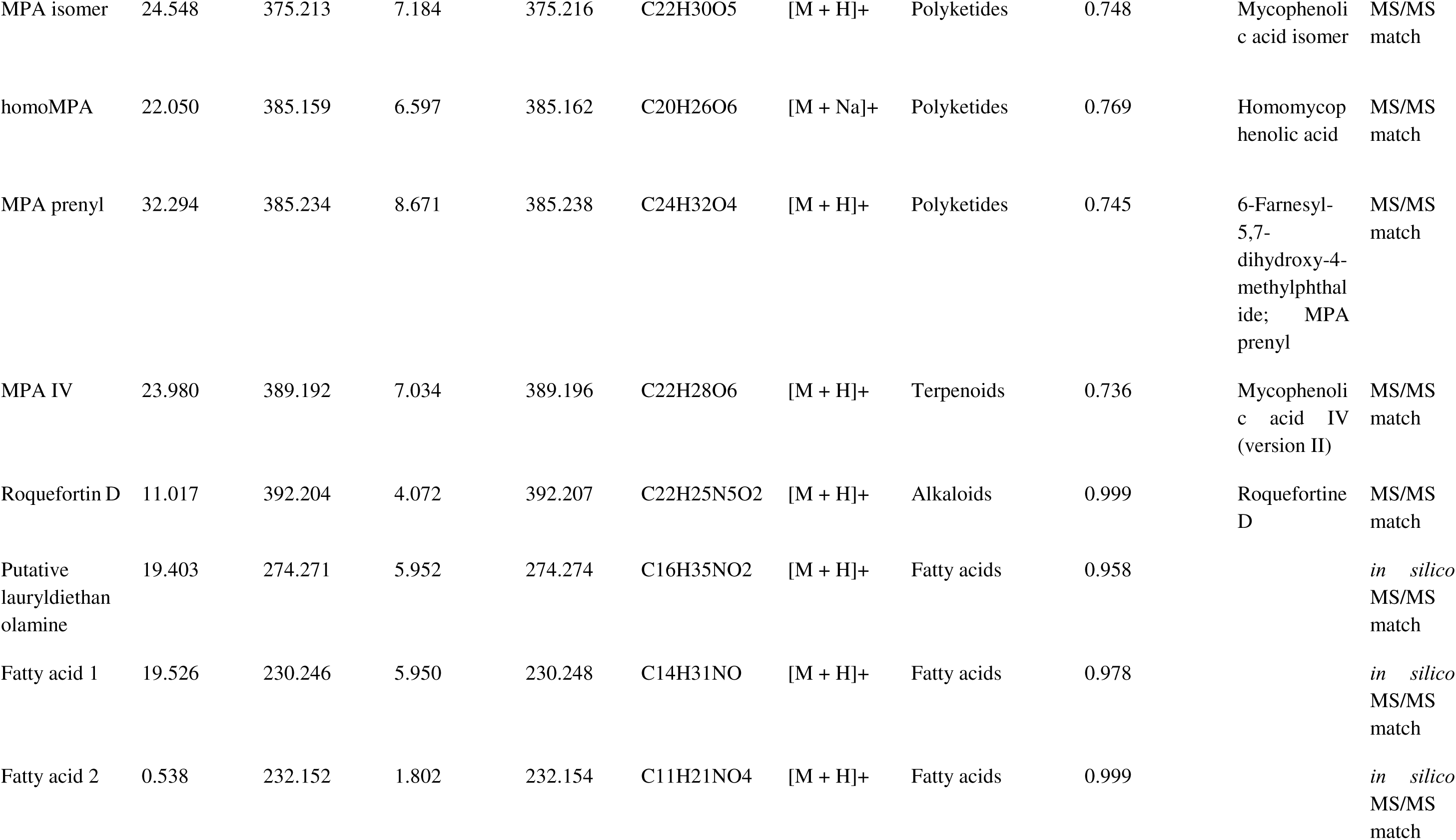
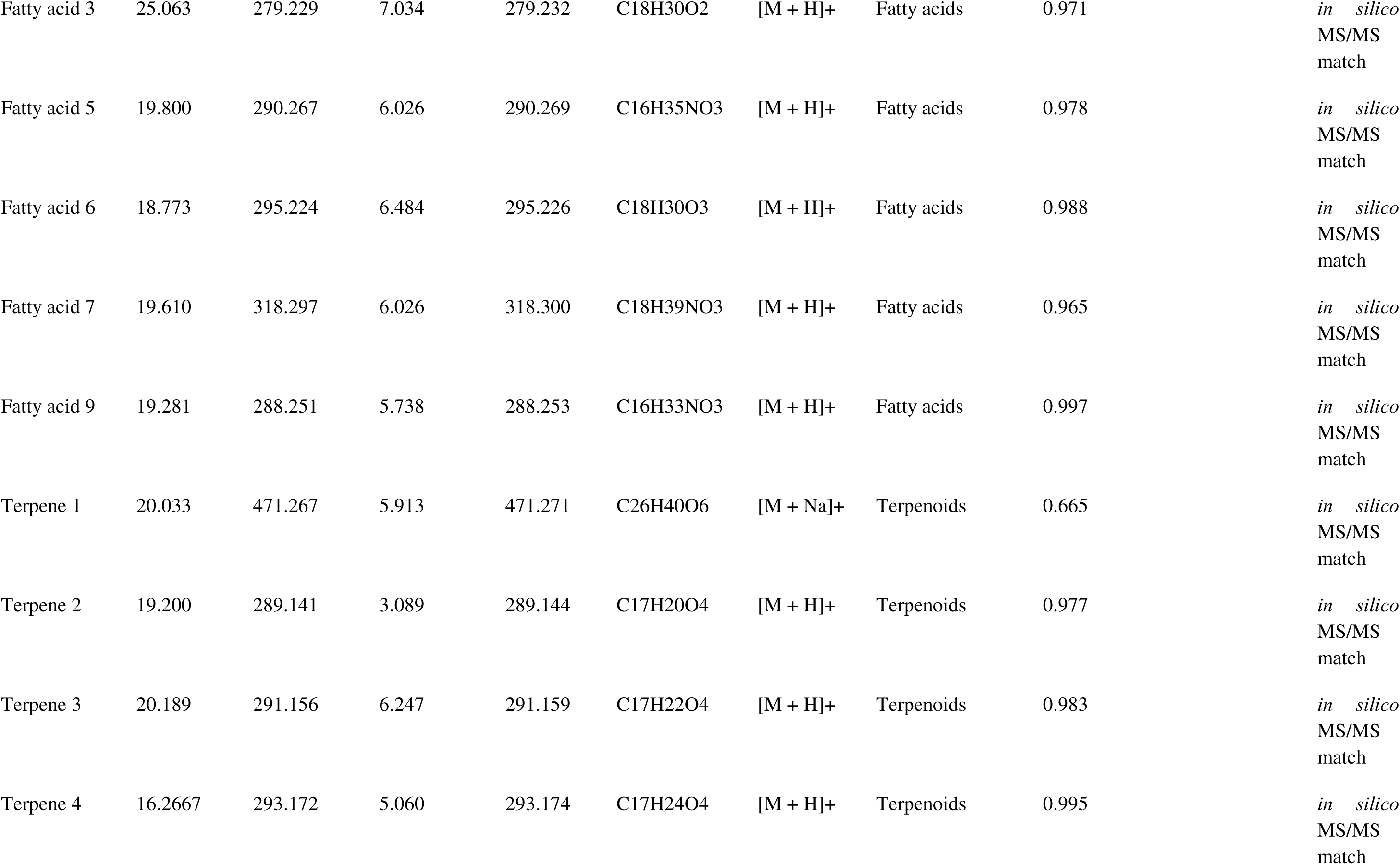
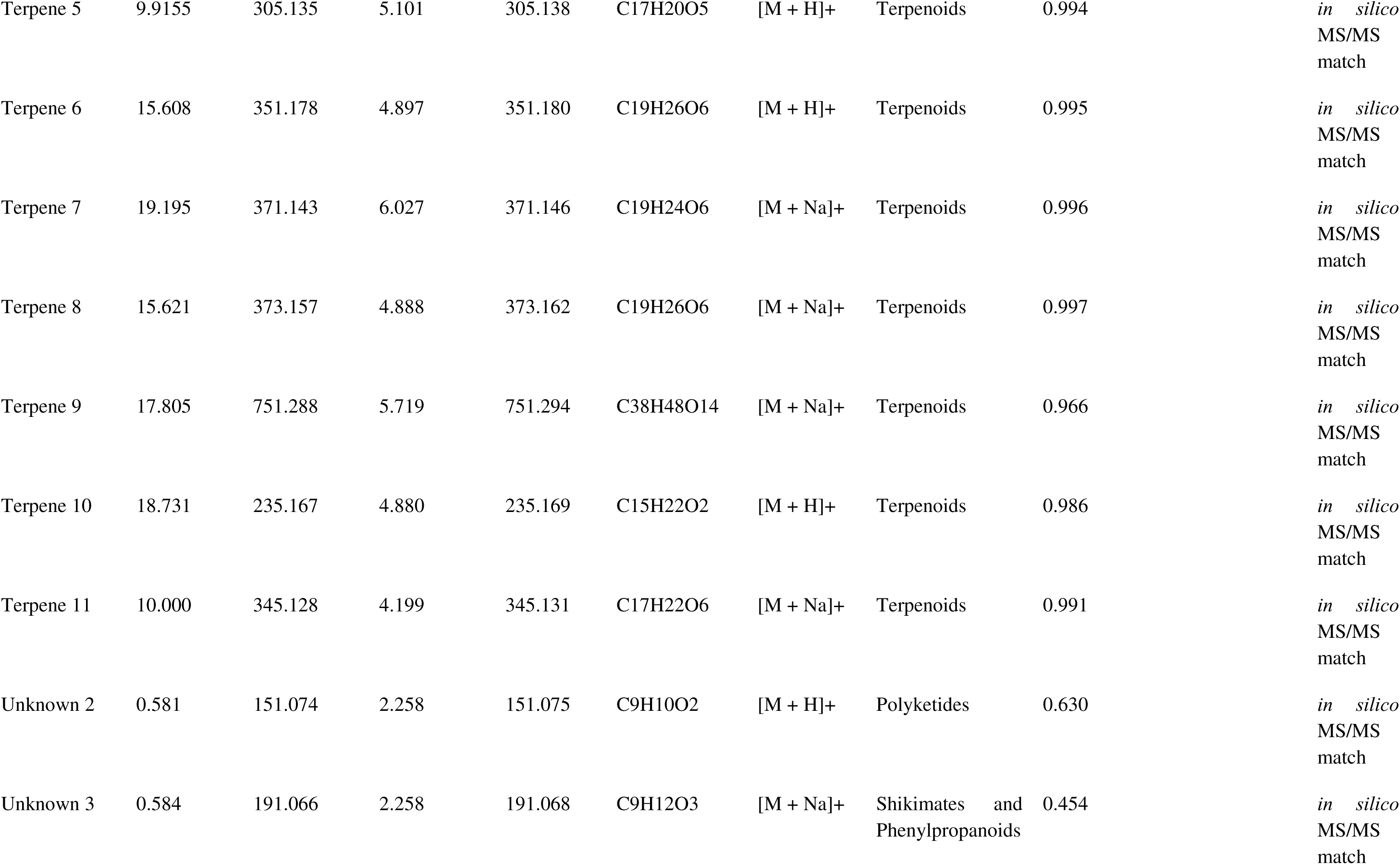
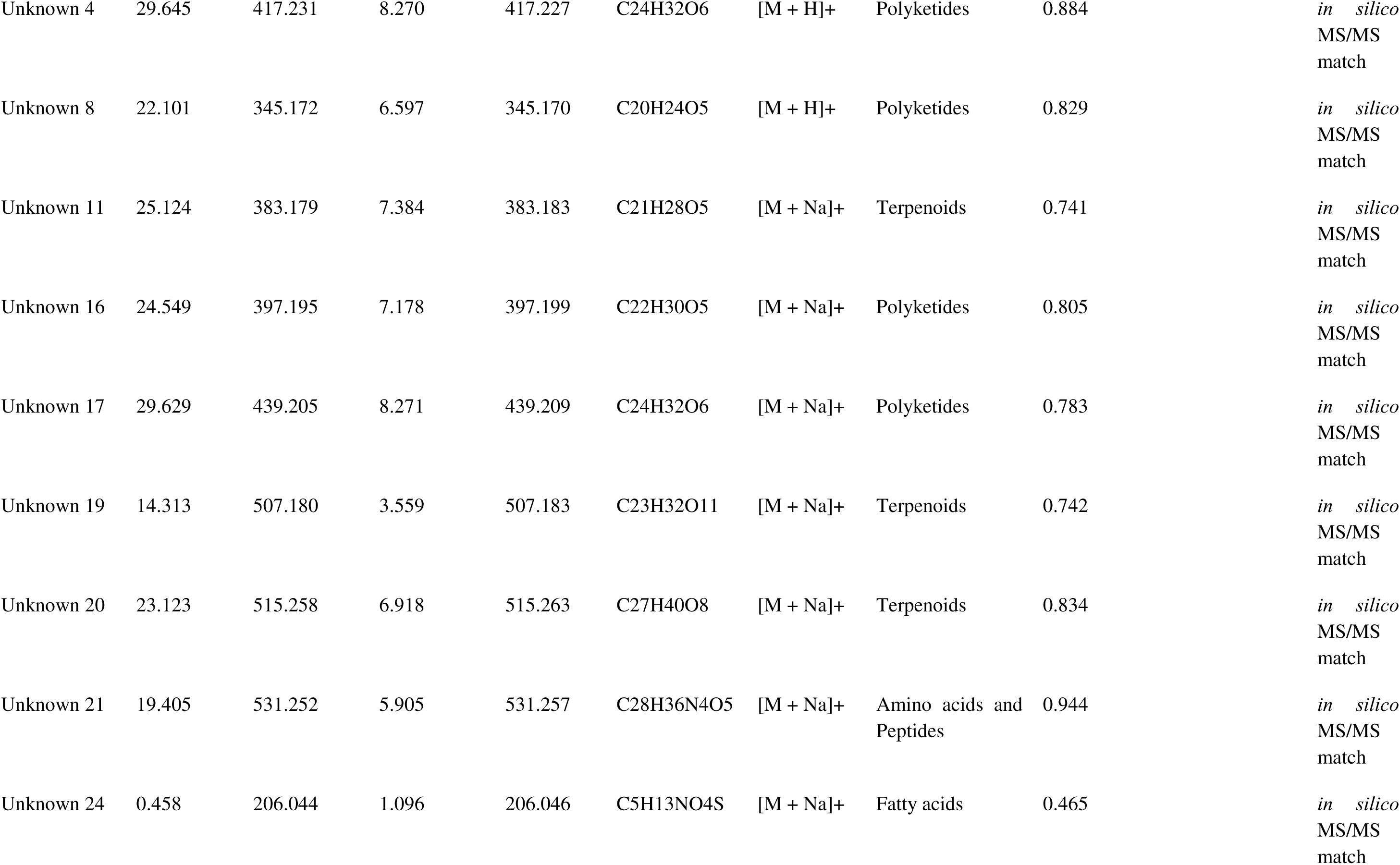
Untargeted metabolite characteristics : Retention times and masses in quantification runs in HPLC-QTOF and identification runs with UHPLC-QTOF, Sirius prediction (*in silico* MS/MS match) and identification against local databases specific to fungus metabolites (MS/MS match).

### Distinct differences in metabolite profiles between cheese and non-cheese populations

The metabolite profiles were indeed strikingly different between the cheese and non-cheese populations. The most toxic mycotoxin produced by *P. roqueforti,* PR toxin, was produced only by non-cheese strains (except a slight production in the Termignon population) and in highest quantities by the lumber/spoiled food population (Figure 2A). The non-cheese populations also displayed higher production of the meroterpenoid AND A (Figure 4B), diverse fatty acids, the isofumigaclavine A intermediate, festuclavine (Suppl. Figure 1 B), the unknown compounds 2 and 3 (Suppl. Figure 4 A & B), the terpenes 5 (Suppl. Figure 3 E) and 11 (Figure 3B), as well as other terpenes. In contrast, cheese populations produced very low or even no detectable amounts of these metabolites (Fig 2, Supp Figure 4). In particular, the fatty acids 1, 5, 7, 9 and the putative lauryldiethanolamine (Figures 2A to E) were not detected in cheese strains. The roquefortine C and D alkaloids were in contrast produced in higher quantities in cheese populations than non-cheese populations (Suppl. Figures 1 H & I).

**Figure 2:**
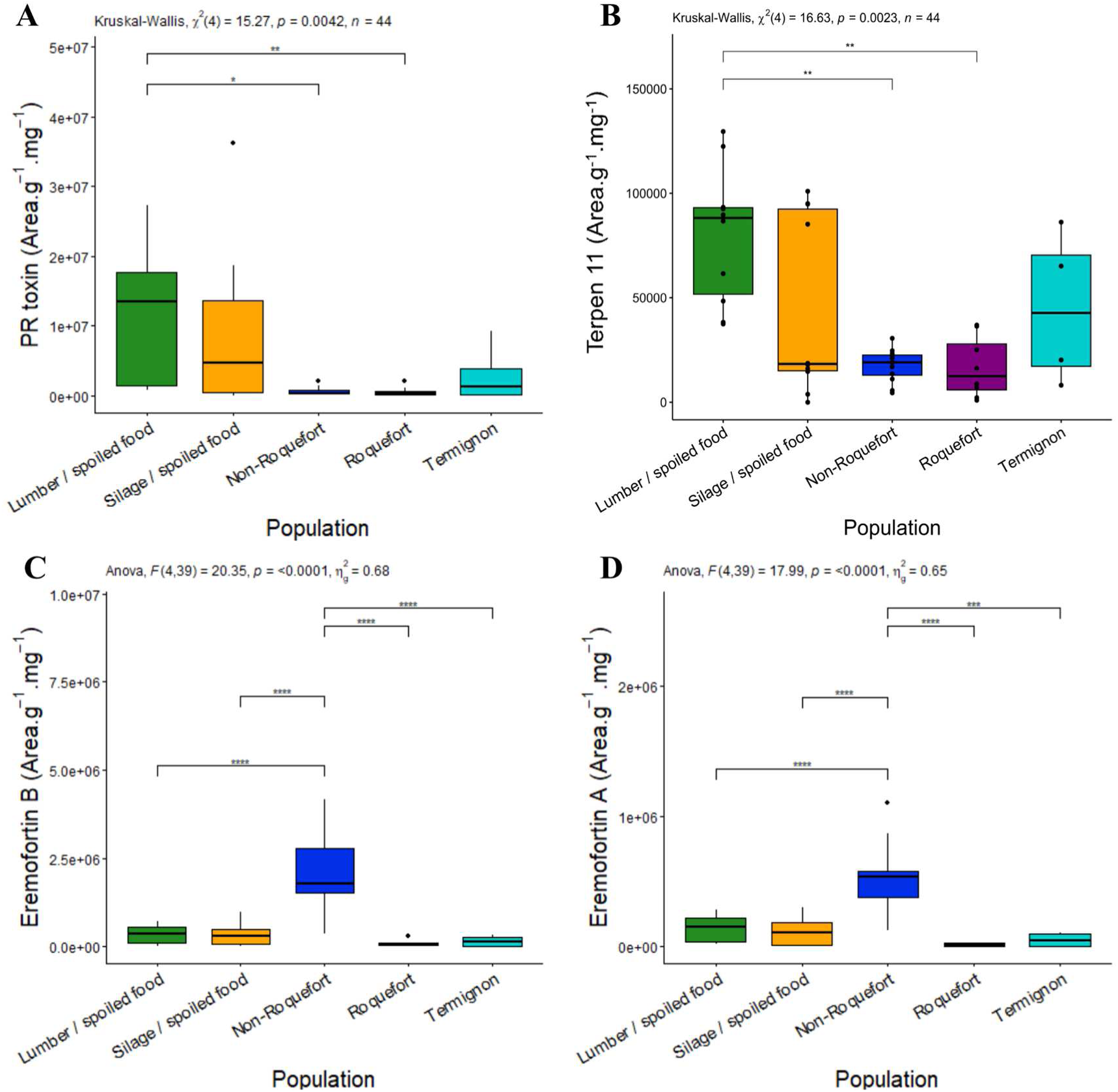
Production level of PR-toxin (A), terpene 11 (putative eremofortin C) (B), eremofortins A (C) and B (D) PR toxin intermediates, across the five *Penicillium roqueforti* populations. Production level is expressed as the surface of the peak area of the targeted metabolite per extract matrix mass and mycelium mass. The same colour code is used as in the other figures: green for the lumber/spoiled food population, orange for the silage/spoiled food population, dark blue for the non-Roquefort cheese population, purple for the Roquefort cheese population and light blue for the Termignon cheese population. The results of the stat test for a population effect is given at the top of each panel. Pairwise significant differences are indicated by asterisks. The boxplots represent the median (centre line), the first quartile and third quartile (box bounds), the maximum and minimum excluding outlier points (whiskers), points being the outliers, *i.e.* with values either below the first quartile minus 1.5 fold the interquartile range or above the third quartile plus 1.5 fold the interquartile range.

**Figure 3:**
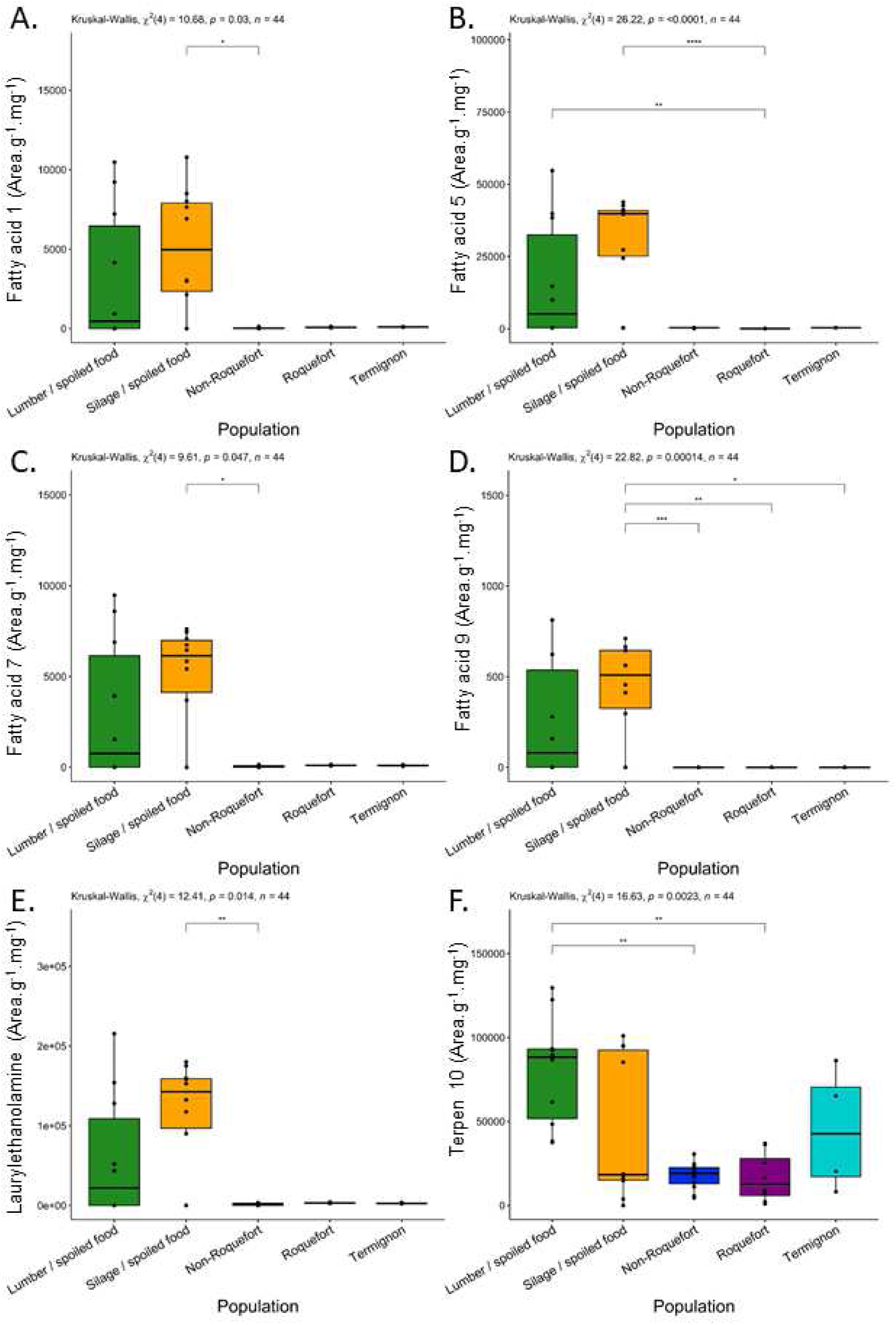
Production levels of the fatty acids 1 (A), 5 (B), 7 (C), 9 (D), laurylethanolamine (E), and terpene 10 (F) across the five *Penicillium roqueforti* populations. Production level is expressed as the surface of the peak area of the metabolite. The different populations were colour-coded as follows: green for the lumber/spoiled food population, orange for the silage/spoiled food population, dark blue for the non-Roquefort cheese population, purple for the Roquefort cheese population and light blue for the Termignon cheese population. The results of the stat test for a population effect is given at the top of each panel. Pairwise significant differences are indicated by asterisks. The boxplots represent the median (centre line), the first quartile and third quartile (box bounds), the maximum and minimum excluding outlier points (whiskers), points being the outliers, *i.e.* with values either below the first quartile minus 1.5 fold the interquartile range or above the third quartile plus 1.5 fold the interquartile range.

**Figure 4:**
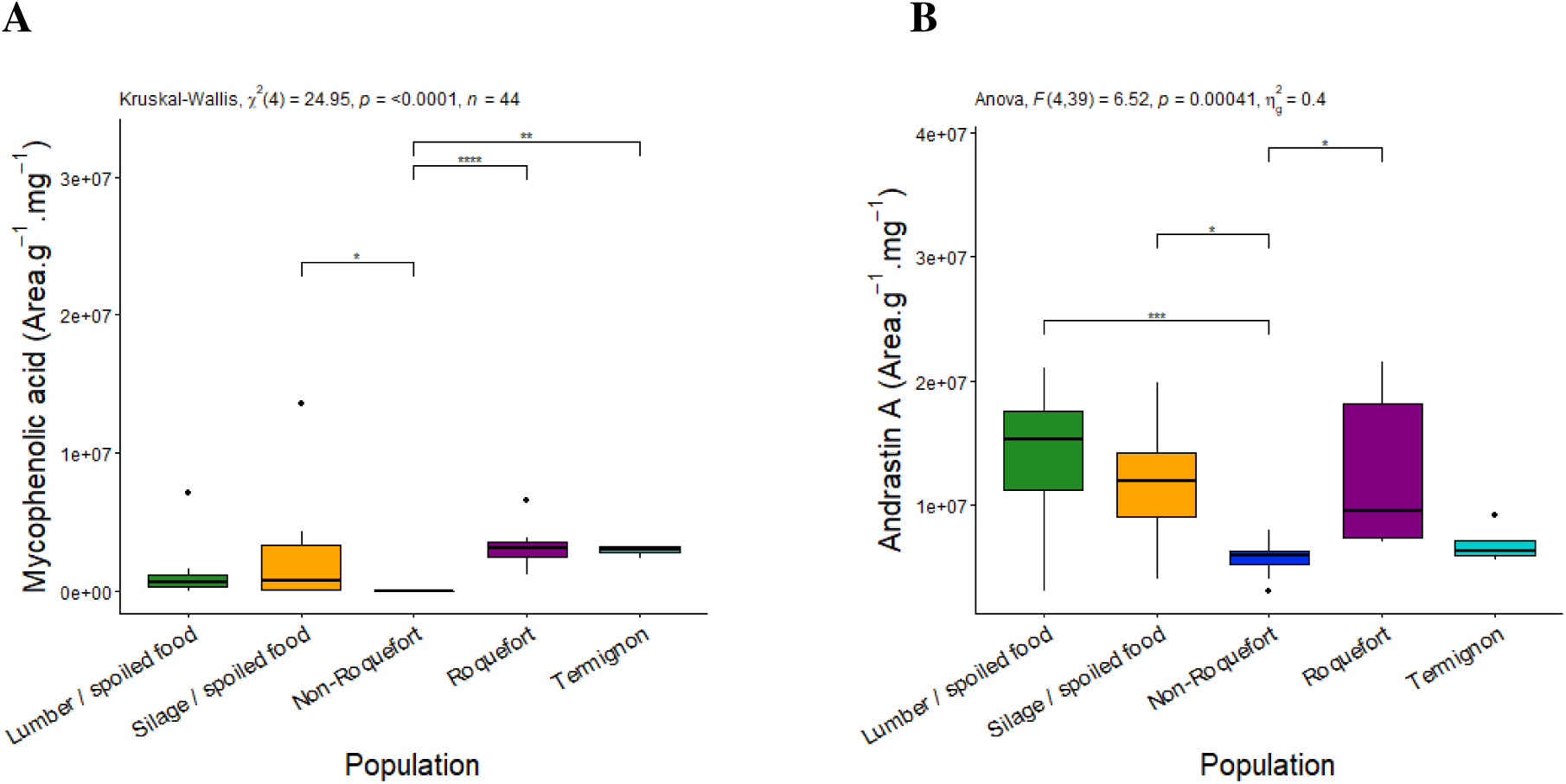
Production level of mycophenolic acid (A) and andrastin A (B) among the five *Penicillium roqueforti* populations. Production level is expressed as the surface of the peak area of the targeted metabolite per extract matrix mass and mycelium mass. The different populations were colour-coded as follows: green for the lumber/spoiled food population, orange for the silage/spoiled food population, dark blue for the non-Roquefort cheese population, purple for the Roquefort cheese population and light blue for the Termignon cheese population. The results of the global test for a population effect is given at the top of each panel. Pairwise significant differences are indicated by asterisks. The boxplots represent the median (centre line), the first quartile and third quartile (box bounds), the maximum and minimum excluding outlier points (whiskers), points being the outliers, *i.e.* with values either below the first quartile minus 1.5 fold the interquartile range or above the third quartile plus 1.5 fold the interquartile range.

### Specific metabolite profiles in each population

The silage/spoiled food population produced the highest levels of two potentially toxic clavines, agroclavine and FUM A (Suppl. Figures 1 A & C), fatty acids 1, 5, 7, 9 and the putative lauryldiethanolamine (Figure 3A). The Roquefort population produced the highest levels of ROQ C & D compared to the other cheese populations (Suppl. Figures 1 H & I). The non-Roquefort population produced the lowest levels of the main mycotoxins across all populations; strains from this population nevertheless produced significantly higher quantities of the PR toxin intermediates, ERE A & B, than the other populations (Figures 2 B & C), suggesting that the PR toxin pathway was partially functional. Only low levels of each targeted metabolite of the PR toxin pathway could be detected in the Roquefort population, a single strain producing quantifiable amounts of a single eremofortin (ERE B for LCP02939; Suppl. Table S3). This suggests that the PR toxin production pathway may be non-functional or down-regulated.

We detected very low quantities of mycophenolic acid across all populations compared to other quantified metabolites, maximal concentrations only reaching 118 ng.g^−1^.mg^−1^. Moreover, 43% of tested strains did not produce any detectable amount of mycophenolic acid, in particular all non-Roquefort strains (Figure 4A). In contrast, all Roquefort and Termignon strains produced detectable MPA levels (Figure 4A). Only Termignon strains produced significant levels of MPA-related metabolites, including identified compounds (MPA isomer, homo MPA, MPA prenyl and MPA IV; Suppl. Figures 1 D, E, F and G) as well as unknown compounds (unknown 4 and 8; Suppl. Figures 4 C & D). The non-Roquefort and Termignon populations produced low levels of AND A, while the Roquefort population produced higher amounts, although in variable quantities (Figure 4B).

Each cheese population also exhibited metabolite specificities in terms of fatty acids and terpenes. The non-Roquefort population produced high levels of fatty acid 2 and terpene 3, and also the highest amounts of the unknown metabolite 24 which harbours a sulfur (Suppl. Figures 2D, 3C and 4K, respectively); sulfur-containing specialised metabolites are often associated with bioactive properties. The Roquefort population produced high levels of unknown metabolites 19, 20 and 21 (Suppl. Figure 4H, I and J), while the Termignon strains produced at higher levels the unknown metabolites 4, 8, 11 and 16 (Suppl. Figure 4 C, D, H, I and J).

### Genetic specificities explain MPA and PR-toxin production differences

The genomes of the 44 studied strains were analysed to compare their biosynthetic gene clusters in order to understand the observed differences in terms of PR toxin and MPA production. For the MPA biosynthetic gene cluster, genomic comparisons showed that all strains from the non-Roquefort population, producing no MPA, exhibited a 174 bp deletion in the *mpaC* gene (Figure 5A). The deletion is situated in the lipase/esterase domain of the MpaC enzyme, which is the key polyketide synthase enzyme of this cluster (Gillot et al. 2017a). Indeed, this nonreducing polyketide synthase catalyses the synthesis of the first reaction intermediate, 5-methylorsellinic acid (5-MOA) from acetyl-CoA, 3 malonyl-CoA and S-adenosylmethionine (Regueira et al. 2011). This deletion in the 3’ region of the gene introduces a frameshift in the translated protein, leading to a truncated protein (2477 aa) compared to the normal-sized protein (2491 aa) of the MpaC sequence found in the other populations (Figure 5A). Beyond the non-Roquefort strains, seven others did not produce quantifiable levels of MPA (three out of ten strains from the lumber/spoiled food population and four out of ten strains from the silage/spoiled food populations, respectively) but no *mpaC* gene deletion or other modifications affecting the gene cluster were detected in their genomes.

**Figure 5:**
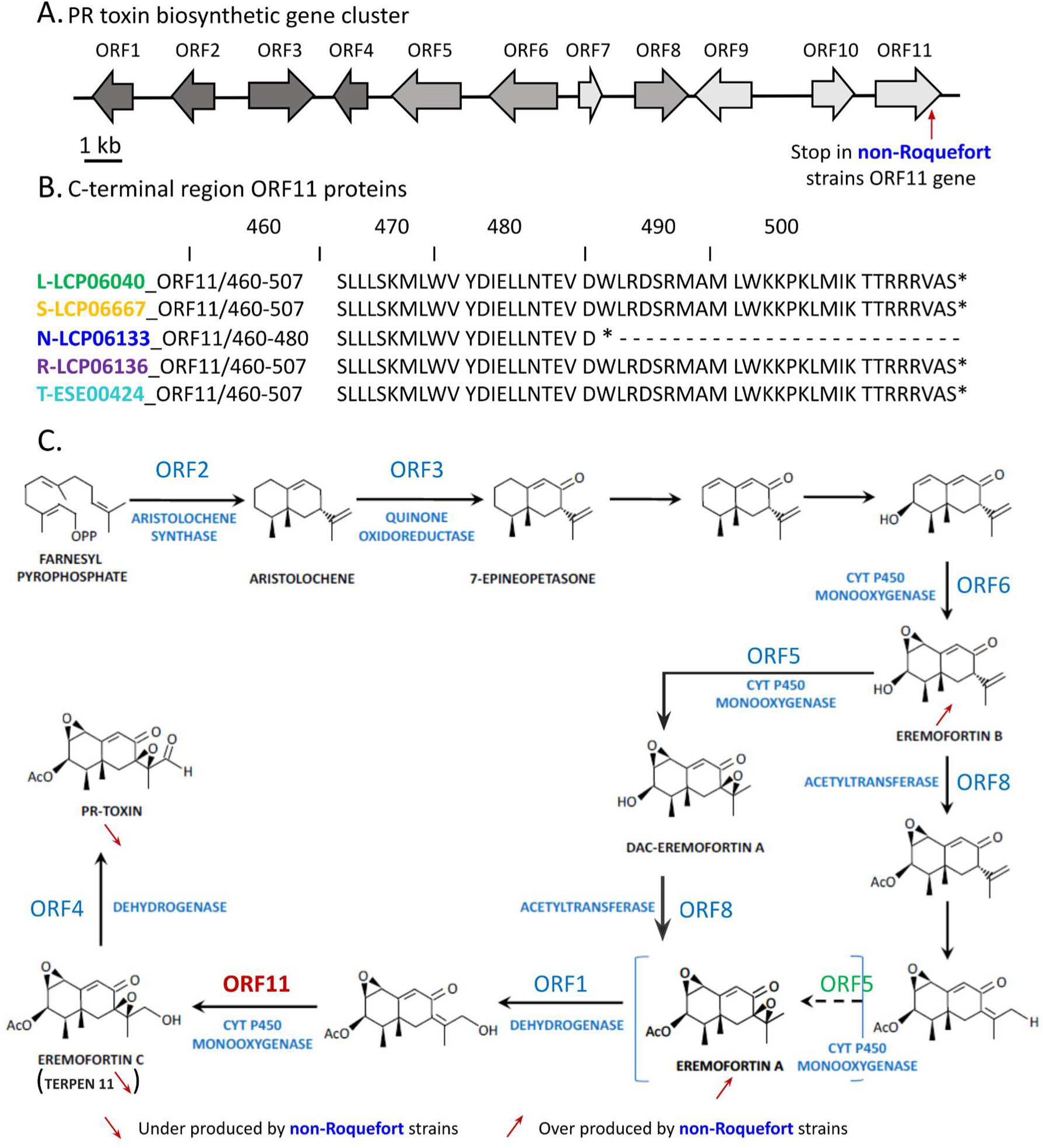
(A) PR toxin biosynthetic gene cluster in *Penicillium roqueforti* as described in Hidalgo et al (2017). Gene silenced in Hidalgo et al (2014) are in dark grey, gene silenced in Hidalgo et al (2017) and untargeted genes are in light grey. (B) C-terminal region of ORF11 protein in strain representing sequenced in their population, “lumber/ spoiled food” in green, “silage/spoiled food” in orange, “non-Roquefort” in blue, “Roquefort” in purple, and “Termignon” in light blue. (C) Proposed position the intervention of ORF11 (red) protein in the PR-toxin production pathway. Pathway figure adapted from Chàvez et al (2023). Position of ORF 5 is not the one proposed in Hidalgo et al (2017) but coherent with an under production of eremofortin A and PR toxin, and an overproduction of eremofortin B observed when the gene is silenced

The genomic comparison of the PR toxin biosynthetic gene clusters revealed that all studied non-Roquefort strains presented a G-to-A substitution in position 1440 of ORF 11, which codes for a cytochrome P450 monooxygenase (Figure 5B). This substitution introduces a premature stop codon instead of a tryptophan codon, resulting in a truncated version of the enzyme, with 27 aa missing in the 3’ region, leading to a protein of 480 aa instead of 507 aa. The truncation in non-Roquefort strains, together with their lack of PR toxin production and the accumulation of the ERE A & B intermediates, suggests that the truncated enzyme is not fully functional. In the Roquefort population, no PR toxin was produced and ERE A & B did not accumulate, but the PR toxin biosynthetic gene cluster displayed no differences with the functional gene clusters in the other *P. roqueforti* populations (Figure 5B). The lack of PR toxin production and of its intermediates may thus be due to *trans*-down-regulation.

A premature stop codon was also observed in the *ifgB* gene of 10 strains from silage, lumber/spoiled food and Termignon population, resulting in a truncated protein of 338 instead of 340 amino acids. However, no lower production of FUM A was observed for the concerned strains compared to others. Also, a frameshift was observed in the *ifgI* gene of the LCP06040 from lumber/spoiled food population, resulting in a truncated protein of 304 instead of 384 amino acids. Interestingly, LCP06040 was the only *P. roqueforti* strain that did not produce FUM A. Concerning the roquefortine C biosynthesis gene cluster, the 5’ region of the *rds* gene has a deletion leading to the absence of a start codon in UBOCC-A-118017 and UBOCC-A-118018 from the silage population as well as a frameshift due to a deletion in the strain LCP02939 from the Roquefort population. The three strains are the only strains which did not produce any roquefortine C. However, none of these four mutations were linked to a given population. For AND A, while allelic variations were observed in some strains, these modifications did not impact the gene cluster and were not associated with a given population.

## 4. Discussion

In this study, we compared metabolite production patterns between strains belonging to the five known *P. roqueforti* populations. We found that the domesticated Roquefort and non-Roquefort cheese populations produced fewer metabolites and were less mycotoxinogenic than their non-cheese counterparts. These findings provide a more thorough understanding on the population divergence and domestication history of *P. roqueforti.* The lower toxin production levels in domesticated populations may be due to selection for healthier cheeses, or to relaxed selection if toxins are not useful any more in the cheese environment compared to wild or other anthropized environments (Lu et al. 2006; Rokas 2009).

To understand adaptation and specialisation in distinct ecological niches, comparing phenotypes of strains associated with different environments is essential to identify adaptive traits. In fungi, these approaches include estimating the growth cardinal values (*e.g.* temperature, pH or *a_w_*), ability to use different substrates and resistance to toxic compounds as recently done for example in *P. roqueforti* (Dumas et al. 2020; Crequer et al. 2023) and in the rice blast fungus *Pyricularia oryzae* (Thierry et al. 2022). Comparisons of metabolite production profiles, especially specialised metabolites, are of major interest in this context as metabolites can be involved in a wide range of biotic interactions and abiotic responses, which can significantly impact fitness and confer competitive advantages.

In this study, we analysed 44 strains from the five known *P. roqueforti* populations (Dumas et al. 2020; Crequer et al. 2023), using a metabolomics approach to compare their metabolite profiles after growth on YES medium and genomic comparison of their metabolite biosynthetic gene clusters. Beyond the seven targeted specialised metabolites, which included the main known *P. roqueforti* mycotoxins, 40 other fungal metabolites were identified (Table 1). The metabolite production profiles were different between the five *P. roqueforti* populations, and in particular between the cheese and non-cheese populations, which we could explain for some extrolites by deletions in genes involved in their biosynthesis in the non-Roquefort population. A major finding relates to the PR toxin, the most toxic known *P. roqueforti* metabolite, with significant differences of production levels between the non-cheese and cheese populations. The non-cheese populations produced, on average, higher concentrations of PR toxin, especially the Lumber/spoiled food population, while the non-Roquefort and Roquefort domesticated populations did not produce any detectable quantities. The Termignon population had an intermediate profile, as previously reported for various growth parameters and carbon source usage (Crequer et al. 2023). Such intermediate metabolite production levels are consistent with the hypothesis that the Termignon population represents descendants of an ancestral domesticated population, displaying traits resulting from domestication before the strong selection imposed in recent years by process industrialisation, thus corresponding to a protracted domestication process (i.e. a slow process occurring across hundreds or thousands of years), as reported in several crops (Allaby et al. 2008; Fuller et al. 2012). Regarding the two populations used for cheese inoculation, non-Roquefort and Roquefort, our results are of particular interest for food safety and human health, as these populations did not produced PR toxin which is the most toxic *P. roqueforti* mycotoxin (Pedrosa and Griessler 2010; Hymery et al. 2017).

On the other hand, *P. roqueforti* is one of the most common post-harvest fungal contaminants in silages (Gallo et al. 2015). Its ability to colonise this substrate, and produce there PR toxin, has been associated with cattle intoxication, with symptoms such as loss of appetite, cessation of rumen activity, gastroenteritis, haemorrhage and even death (Veselý et al. 1981; Nielsen et al. 2006). It was considered so far that the *P. roqueforti* strains used as ripening cheese cultures had the intrinsic ability to produce PR toxin (Dubey et al. 2018) and that the absence of this mycotoxin in cheeses was due to its instability and presumed degradation into various less toxic molecules, i.e. PR imine and PR amide and PR acid (Chang et al. 1993, 1996). While the latter hypothesis may still be valid, our results provide a new and robust explanation, as none of the studied non-Roquefort and Roquefort strains, used for blue cheese production, produced quantifiable levels of PR toxin, even in YES medium known to be favourable for secondary metabolite production. Cheeses made with potential PR-toxin producer strains, such as the Termignon population, may nevertheless contain little of this toxin in cheeses as it is unstable in this matrix. Indeed, PR toxin was shown to be degraded in PR amine in cheese (Siemens and Zawistowski 1993). The instability of PR toxin in cheese may imply that the loss of its production ability in cheese strains may be more due to relaxed selection than to selection against its production in cheeses.

Diversity in metabolite synthesis, both qualitatively and quantitatively, may arise from divergent selection across ecological niches. Different metabolite profiles across differentiated fungal populations or lineages of a given species have been reported in other fungi. For example, among the four lineages identified in *Fusarium graminearum* isolated from maize (Lee et al. 2012), most isolates from lineages 2 and 6 produced the trichothecene group B mycotoxin nivalenol (NIV), while all isolates from lineages 3 and 7 produced deoxynivalenol (DON), another major trichothecene B *Fusarium* mycotoxin, that is a virulence factor in wheat and toxic for human and animal health. In *Fusarium asiaticum* isolated from Chinese rice and wheat, producer strains of 3-acetyldeoxynivalenol (an acetylated form of DON) were ubiquitous in wheat while NIV-producers were more prevalent in rice, the trichothecene chemotypes also varying across regions (Yang et al. 2018). In *Aspergillus flavus,* the production of aflatoxin B1 (AFB1), a potent cancerogenic mycotoxin regulated in the food chain, was significantly higher for soil isolates than for corn kernel ones (Sweany et al. 2011).

To further understand the differences in metabolite production between *P. roqueforti* populations, we focused on determining the genetic basis for two main differences between cheese and non-cheese populations, i.e. the production of PR toxin and MPA. The inability to produce PR toxin by non-Roquefort strains could be attributed to a substitution in ORF 11 of the corresponding biosynthetic gene cluster resulting in a premature stop codon. This finding, combined with the accumulation of both ERE A & B, suggests that ORF 11 likely intervenes in the formation of eremofortin C (ERE C), the final precursor for PR toxin, instead of ORF 5 as previously described (Hidalgo et al. 2017). Our results also pinpointed an unknown metabolite that may correspond to ERE C (terpene 11); this metabolite was not found in the non-Roquefort population extracts either, which reinforces the hypothesis that ORF 11 intervenes in its formation. However, as no commercial ERE C standard was available, we could not fully confirm the identity of terpene 11 nor test the possibility that these isolates can produce PR toxin from ERE C.

The Roquefort population produced no detectable amount of PR toxin, ERE A, ERE B or putative ERE C (*i.e.* terpene 11), and we were unable to determine the genetic basis of this lack of production based on biosynthetic gene cluster comparisons. It seems most likely that the expression of the entire gene cluster might be affected by a regulatory element in *cis* or *trans*. In the DS17690 *P. chrysogenum* strain, downregulation of the PR toxin biosynthetic cluster was due to mutations in the *laeA* and *velA* regulatory genes (Martín 2017). Here, we did not identify any mutations in either of these two genes in the Roquefort strains (data not shown); therefore, the inability to produce PR toxin may be due to identified global regulators (*e.g. pga1, sfk1, pcz1*) involved in the modulation of metabolite production in *P. roqueforti* (Chávez et al. 2023) or to other, unidentified regulators.

None of the non-Roquefort strains produced mycophenolic acid and we could attribute this inability to a deletion in the *mpaC* gene within the corresponding biosynthetic gene cluster. This deletion had been previously reported in *P. roqueforti* strains (Gillot et al. 2017a), and found associated with the presence of the horizontally transferred *CheesyTer* and *Wallaby* regions (Gillot et al. 2017b), that were later found mostly present in the non-Roquefort population (Dumas et al. 2020). The Termignon strains were the highest producers of MPA and MPA-related derivatives, which is of interest for large-scale production of this important pharmaceutical immunosuppressive with antifungal, antibacterial, antiviral, anti-psoriasis and antitumor and anti-graft reject activities (Ammar et al. 2023).

In *P. roqueforti,* other extrolites were produced by all populations, but with still marked differences in production levels for several compounds. For the ROQ C & D alkaloids, the highest producers were found in the cheese populations, especially the Roquefort population. The fact that ROQ C & D production was maintained in domesticated populations raises questions about their ecological role in cheese, given that roquefortines have various bioactive properties. It may also be that there was no selection during domestication against the production of this extrolite with low cytotoxic effects (Fontaine et al. 2016).

We also identified differences between *P. roqueforti* populations for the production of clavines, *e.g.* FUM A, festuclavine (a FUM A intermediate) and agroclavine. The production of festuclavine and agroclavine by *P. roqueforti* had previously been reported but not compared between populations (Ohmomo et al. 1975; Nielsen et al. 2006). We found the highest quantities of these compounds in the non-cheese populations, and also in the Roquefort population. Similar results were also observed for andrastin A, a potential natural anti-cancer compound, thus raising the question of the ecological role of this molecule, especially in cheese populations.

Numerous other untargeted molecules were observed and corresponded to terpenoids, fatty acids (including fatty acid amides) or unknown molecules. These molecules might correspond to metabolites recently described in *P. roqueforti,* such as annullatins (Xiang et al. 2022), eremophilane and guaiane sesquiterpenes (Mo et al. 2023) or sesterterpenoids (Wang et al. 2018, 2020, 2021), their role and biological activity being still unknown. Further efforts are required to refine their identification and understand their function. Several molecules were specific to some populations, *e.g.* unknown compounds 2 and 3, and terpene 11 specific to the lumber population, fatty acid 5 and putative lauryldiethanolamine specific to the silage/spoiled food population, terpene 3 specific to the non-Roquefort population, unknown compounds 20 and 21 specific to Roquefort population, and unknown compounds 8 and 11 specific to the Termignon population; they could thus be of clear interest as potential population biomarkers and for being involved in niche specialisation.

The non-Roquefort population, which displays the strongest domestication syndrome and the most severe genetic bottleneck (Dumas et al. 2020; Crequer et al. 2023), exhibited also the most distinctive metabolite profile. The non-Roquefort strains produced the lowest amounts of metabolites for both identified compounds (including mycotoxins) or unidentified compounds. Such toxin loss represents a convergence in domesticated fungal populations, and may result from a neutral degeneration of unused traits or a selection against toxin production by humans. Fungal metabolites are known to be used for microbial “chemical warfare”, *e.g.* for fungal invasion in plants and/or microbial competition, so they might not be required to the same extent in a rich medium with readily available nutrients and with an inoculation advantage for matrix colonisation. Other unused traits have been reported to degenerate in domesticated fungi by relaxed selection, such as the ability of carbohydrate use and of sexual reproduction (Ropars et al. 2016; Ropars and Giraud 2022). In the present study, genomic comparisons allowed identifying the mutations likely causing loss of production of the PR toxin and MPA in the non-Roquefort population. *P. roqueforti* was also shown to harbour a non-functional gene cluster for the toxic mycotoxin patulin ((Nielsen et al. 2017).

If there was selection of strains unable to produce harmful compounds in the Non-Roquefort and Roquefort populations, this raises fascinating questions concerning the empiric selection process used as, at the time, neither genetic nor metabolomic data were available. Cheese producers inoculated spores into breads to cultivate *P. roqueforti* strains (bread crusts being burnt to avoid external contaminants), and they likely have inoculated bread using spores from the best cheeses, in particular those causing no health issue.

The loss of mycotoxinogenesis can indeed be the result of domestication events, as humans have often selected fungal strains unable to produce harmful toxins for use in food. The best known example is *Aspergillus oryzae,* a domesticated species used to ferment Asian food products derived from its mycotoxin-producing wild relative, *Aspergillus flavus* (Barbesgaard et al. 1992). In *A. oryzae*, several mutations were reported in the aflatoxin biosynthetic gene cluster, in particular an approximately 40 kb deletion in the genomic region between the *norB* and *norA* genes (Chang et al. 2005), mutations in the *aflR* promoter, a nearly 250 bp deletion in the *aflT* coding region, a frameshift mutation in the *norA* coding region and multiple nonsynonymous mutations in the *verA* coding region (Tominaga et al. 2006). Down-regulation of another mycotoxin, cyclopiazonic acid (CPA), also occurs in *A. oryzae* (Gibbons et al. 2012). Another example is the domesticated fungus *Aspergillus sojae,* a species considered to be derived from *Aspergillus parasiticus* (Chang and Hua 2023). In *A. sojae,* the inability to produce aflatoxin is the result of a termination point mutation in the *aflR* regulatory gene as well as a premature stop codon in the *pksA* gene leading to a truncated version of the polyketide synthase enzyme (Chang et al. 2007). Another example, among *Penicillium* species, is *P. camemberti*, domesticated for cheese making. In this species, two different lineages display very contrasted extrolite production profiles: *P. camemberti* var. *camemberti* strains produce high levels of cyclopiazonic acid (CPA) on YES medium while *P. camemberti* var. *caseifulvum* does not. This was shown to be due to a 2 bp deletion in the *cpaA* gene, inducing a frameshift, thus modifying the polyketide synthase/non-ribosomal peptide synthase enzyme responsible for the first step of the CPA biosynthetic pathway (Ropars et al. 2020).

## 5. Conclusion

To conclude, a dual targeted and untargeted metabolomics approach was used to compare *P. roqueforti* metabolite profiles. Distinct profiles were identified across the five *P. roqueforti* populations which is likely due to ecological specialisation and human selection. Indeed, the two domesticated populations used to inoculate blue cheeses no longer produce PR toxin, the most toxic *P. roqueforti* mycotoxin, while the Termignon population strains produce low levels. In contrast, the non-cheese populations (Lumber/spoiled food and Silage) maintained their PR toxin production which indicates that this mycotoxin likely plays an important ecological role in these more complex and harsh environments where microbial competition and natural colonisation occurs, although its precise role remains unknown. The metabolite diversity and quantity profile is unique to each *P. roqueforti* population and likely provides specific advantages to thrive in their respective complex domesticated or wild environments. Overall, this study provides new findings supporting that fungal metabolite profiles are a result of adaptation to environmental conditions (*i.e.* niche specialisation) and that domestication leads to hypotoxigenic populations.

## Statements & Declarations

### Funding

This study was funded by the ANR-19-CE20-0002-02 Fungadapt (Agence Nationale de la Recherche) grant.

### Competing Interests

The authors have no relevant financial or non-financial interests to disclose.

### Author contributions

Obtained funding: ECo, TG; Designed the work: MC, ECo, JLJ. Performed targeted analyses: GC, ECr, MC; Performed and analysed untargeted analyses: MC, ECr, YC, JF; Metabolomics data analyses MC, ECr, YC, JF; Global data set comparisons ECr; Genomes: TG, ECr, RRdlV; biosynthetic gene cluster analyses: ECr, ECo, JLJ, MC, RRdlV; Statistical analyses: ECr; Writing original draft preparation: ECr, ECo, MC; Editing and proofreading: JLJ and TG with contributions by all authors; Supervision: MC, ECo, JLJ, TG.

### Data Availability

The genome datasets generated during and/or analysed during the current study are available in GenBank via the accession numbers provided. All other datasets generated during and/or analysed during the current study are available from the corresponding author on reasonable request.”

## Acknowledgments

This study was funded by the ANR-19-CE20-0002-02 Fungadapt (ANR: National French Research Agency, “Agence Nationale de la Recherche”) grant. The authors are thankful to Thibault Caron and Jean-Philippe Vernadet for useful advice on bioinformatic analyses.

## Notes

### Competing Interest Statement

The authors have declared no competing interest.

