## Supplemental Tables S1, S2; Supplemental figures 1, 2, 3, 4 for "Different metabolite profiles across *Penicillium roqueforti* populations associated with ecological niche specialisation and domestication"

| **Table S1:** ***Penicillium roqueforti*** **strains used for metabolite profiling, with their IDs, assigned genetic population, sampling origin and date, genome accession number when available and associated reference.** | | | | | | | |
|---|---|---|---|---|---|---|---|
| **Strain code** | **Short ID** | **Population** | **Sampling matrix** | **Sampling country** | **Sampling date** | **Genome accession number** | **Reference** |
| LCP06040 | L1 | Lumber / spoiled food | Drying Wood | France | 1992 | ERS162892 1 | Dumas et al. 2020 |
| LCP05419 | L3 | Lumber / spoiled food | Inner fridge wall | France | Unknown | ERS162891 7 | Dumas et al. 2020 |
| LCP06039 | L4 | Lumber / spoiled food | Fruit Compote | URSS | 1973 | ERS162892 0 | Dumas et al. 2020 |
| LCP04111 | L5 | Lumber / spoiled food | Wood | France | Unknown | ERS162891 4 | Dumas et al. 2020 |
| LCP06060 | L6 | Lumber / spoiled food | Sulphite Liquor | Canada | Before 2014 | ERS162892 4 | Dumas et al. 2020 |
| LCP06064 | L8 | Lumber / spoiled food | Mouldy baker's yeast | Denmark | Unknown | ERS162892 5 | Dumas et al. 2020 |
| LCP06037 | L9 | Lumber / spoiled food | Drying Wood | France | Unknown | ERS162891 9 | Dumas et al. 2020 |
| UBOCC-A-117107 | L10 | Lumber / spoiled food | Apple | Kazakhstan | 1974 | PRJNA9335 71 | Crequer et al. 2023 |
| UBOCC-A-117110 | L11 | Lumber / spoiled food | Soil | Russia | 1982 | PRJNA9335 71 | Crequer et al. 2023 |
| UBOCC-A-117111 | L12 | Lumber / spoiled food | Apple | Kazakhstan | 2007 | PRJNA9335 71 | Crequer et al. 2023 |
| UBOCC-A-117112 | L13 | Lumber / spoiled food | Wood | United Kingdom | 2007 | PRJNA9335 71 | Crequer et al. 2023 |
| LCP06138 | N2 | Non-Roquefort | Blue Cheese; Cambozola black label | Germany | Unknown | ERS162893 6 | Dumas et al. 2020 |
| ESE00422 | N4 | Non-Roquefort | Feta cheese | Australia | Before 2014 | PRJNA9335 71 | Crequer et al. 2023 |

| | | | | | | | |
|---|---|---|---|---|---|---|---|
| LCP06134 | N6 | Non-Roquefort | Blue Cheese; Blue Sunshine | USA | Unknown | ERS1628932 | Dumas et al. 2020 |
| LCP06132 | N7 | Non-Roquefort | Blue Cheese; Stilton | England | Unknown | ERS1628930 | Dumas et al. 2020 |
| LCP00146 | N8 | Non-Roquefort | Blue Cheese; Roquefort | France | Unknown | ERS1628909 | Dumas et al. 2020 |
| LCP06129 | N11 | Non-Roquefort | Blue Cheese; Crémeux du Puy | France | Unknown | ERS1628927 | Dumas et al. 2020 |
| FM164 | N12 | Non-Roquefort | Blue Cheese; Gorgonzola | Unknown | Unknown | HG792015-HG792062 | Cheeseman K, et al. 2014 |
| UBOCC-A-117088 | N13 | Non-Roquefort | Cacao nuts | Unknown (Africa) | Before 2012 | To be added upon acceptance | This article |
| UBOCC-A-117091 | N14 | Non-Roquefort | Subglacial water | Norway | 2006 | PRJNA933571 | Crequer et al. 2023 |
| UBOCC-A-117092 | N15 | Non-Roquefort | Blue Cheese; Bleu du Vercors Sassenage | France | 2016 | PRJNA933571 | Crequer et al. 2023 |
| UBOCC-A-118016 | N16 | Non-Roquefort | Irrigation water | France | 2016 | PRJNA933571 | Crequer et al. 2023 |
| ESE00424 | T1 | Termignon | Blue Cheese; Bleu de Termignon | France | 2018 | PRJNA933571 | Crequer et al. 2023 |
| ESE00426 | T2 | Termignon | Blue Cheese; Bleu de Termignon | France | 2018 | PRJNA933571 | Crequer et al. 2023 |
| ESE00428 | T3 | Termignon | Blue Cheese; Bleu de Termignon | France | 2018 | PRJNA933571 | Crequer et al. 2023 |

| | | | | | | | |
|---|---|---|---|---|---|---|---|
| ESE00685 | T4 | Termignon | Blue Cheese; Bleu de Termignon | France | 2015 | PRJNA933571 | Crequer et al. 2023 |
| ESE00170 | R1 | Roquefort | Peach | Unknown | 2018 | PRJNA933571 | Crequer et al. 2023 |
| LCP06136 | R2 | Roquefort | Blue Cheese; Great Hill Blue | USA | Unknown | ERS1628934 | Dumas et al. 2020 |
| ESE00425 | R3 | Roquefort | Blue Cheese; Bleu de Termignon | France | 2018 | PRJNA933571 | Crequer et al. 2023 |
| LCP02939 | R5 | Roquefort | Brioche under plastic wrap | Unknown | Unknown | ERS1628911 | Dumas et al. 2020 |
| LCP06130 | R6 | Roquefort | Blue Cheese; Gorgonzola | Italy | 2013 | ERS1628928 | Dumas et al. 2020 |
| LCP04157T | R7 | Roquefort | Blue Cheese; Roquefort | USA | 1987 | ERS1628915 | Dumas et al. 2020 |
| UBOCC-A-113022 | R8 | Roquefort | Blue Cheese; Roquefort | France | 2013 | PRJNA933571 | Crequer et al. 2023 |
| UBOCC-A-116029 | R9 | Roquefort | Atmosphere cave | France | 2016 | PRJNA933571 | Crequer et al. 2023 |
| LCP06059 | S1 | Silage / spoiled food | Silage | Netherlands | Unknown | ERS1628923 | Dumas et al. 2020 |
| LCP04180 | S2 | Silage / spoiled food | Strawberry sorbet | France | 1997 | ERS1628916 | Dumas et al. 2020 |
| LCP03969 | S3 | Silage / spoiled food | Fruit Compote | France | 1996 | ERS1628913 | Dumas et al. 2020 |
| LCP06635 | S5 | Silage / spoiled food | Silage | Belgium | 2006 | PRJNA933571 | Crequer et al. 2023 |
| LCP06667 | S7 | Silage / spoiled food | Silage | Belgium | 2007 | PRJNA933571 | Crequer et al. 2023 |
| LCP06679 | S10 | Silage / spoiled food | Silage | Belgium | 2007 | PRJNA933571 | Crequer et al. 2023 |

| | | | | | | | |
|---|---|---|---|---|---|---|---|
| UBOCC-A-117102 | S12 | Silage / spoiled food | Silage | Belgium | 2011 | PRJNA933571 | Crequer et al. 2023 |
| UBOCC-A-117291 | S13 | Silage / spoiled food | Silage | France | 2017 | PRJNA933571 | Crequer et al. 2023 |
| UBOCC-A-118017 | S14 | Silage / spoiled food | Flour | France | 2016 | PRJNA933571 | Crequer et al. 2023 |
| UBOCC-A-118018 | S15 | Silage / spoiled food | Soft bread | France | 2015 | To be added upon acceptance | This article |
| ESE00421 | L2 | Unassigned | Bread | France | 2019 | PRJNA933571 | Crequer et al. 2023 |

**Table S2.** Method peformance characteristics for metabolite quantification in YES medium.

| Compound | RT | Quantifier Ion (Q1) (m/z) | Qualifier Ion (Q2) (m/z) | R2 | LOD (ng.mg-1) | LOQ (ng.mg-1) | LOD (area.mg-1) | LOQ (area.mg-1) | ESI |
|---|---|---|---|---|---|---|---|---|---|
| PR toxin | 17.4753 | 321.1361 | NA | 0.996 | NA | NA | 458857.261116974 | 3125458.11802791 | + |
| Eremofortin A | 16.2030 | 307.1537 | 329.1358 | 0.999 | 188.5822838125 | 571.4614661875 | 10869.3867614338 | 52507.0219496045 | + |
| Eremofortin B | 12.0695 | 249.1482 | 271.1252 | 0.998 | 209.706599125 | 635.4745425 | 32779.9527435967 | 135541.122463209 | + |
| Mycophenolic acid | 17.7789 | 321.1335 | 303.1231 | 0.999 | 128.9628125 | 390.796401875 | 352146.953607505 | 919746.986293351 | + |
| Andrastine A | 22.4684 | 487.2690 | NA | 1.000 | 303.303548125 | 919.1016625 | 7157.80862442707 | 177371.451065421 | + |
| Roquefortine C | 15.2823 | 390.1923 | NA | 0.982 | 1051.288410625 | 3185.7224575 | 1638523.67044636 | 3365946.01417412 | + |
| (Iso-)fumigaclavine A | 3.1594 | 299.1759 | NA | 0.999 | 166.00053625 | 503.031928125 | 528981.401298376 | 1609886.25933455 | + |
| Mycophenolic acid | 13.4512 | 319.1165 | NA | 0.999 | 346.580822 | 1050.244915 | 240305.756875 | 980703.164375 | - |
| Andrastine A | 16.0868 | 485.2545 | NA | 0.960 | 3100.244051875 | 9394.67894375 | 5813301.9125 | 10408340.94375 | - |

RT: Retention time; R2: determination coefficient; DL: Deterction limit; QL: Quantification limit; ESI: Electrospray ionization; NA: not applicable

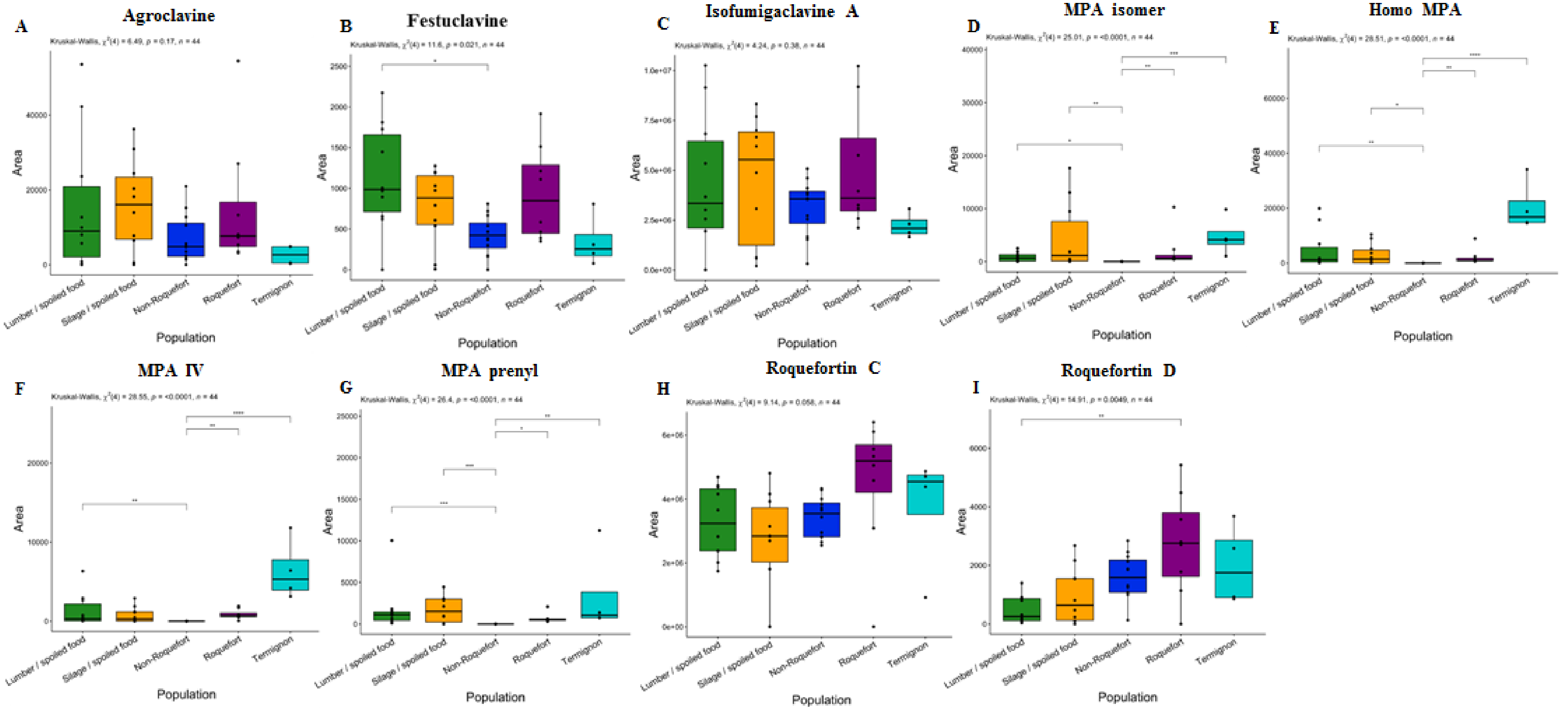


**Supplementary Figure 1: Production level of agroclavine (A), festuclavine A (B), Isofumigaclavine A (C), mycophenolic acid (MPA) isomer (D), homo-MPA (E), MPA-IV (F), -MPA-prenyl (G), roquefortine C and (H) roquefortine D (I) among the five *Penicillium roqueforti* populations.** Production level is expressed as the surface of the peak area of the targeted metabolite per extract matrix mass and mycelium mass. The different populations were colour-coded as follows: green for the lumber/spoiled food population, orange for the silage/spoiled food population, dark blue for the non-Roquefort cheese population, purple for the Roquefort cheese population and light blue for the Termignon cheese population. The results of the global test for a population effect is given at the top of each panel. Pairwise significant differences are indicated by asterisks. The boxplots represent the median (centre line), the first quartile and third quartile (box bounds), the maximum and minimum excluding outlier points (whiskers), points being the outliers, *i.e.* with values either below the first quartile minus 1.5 fold the interquartile range or above the third quartile plus 1.5 fold the interquartile range.

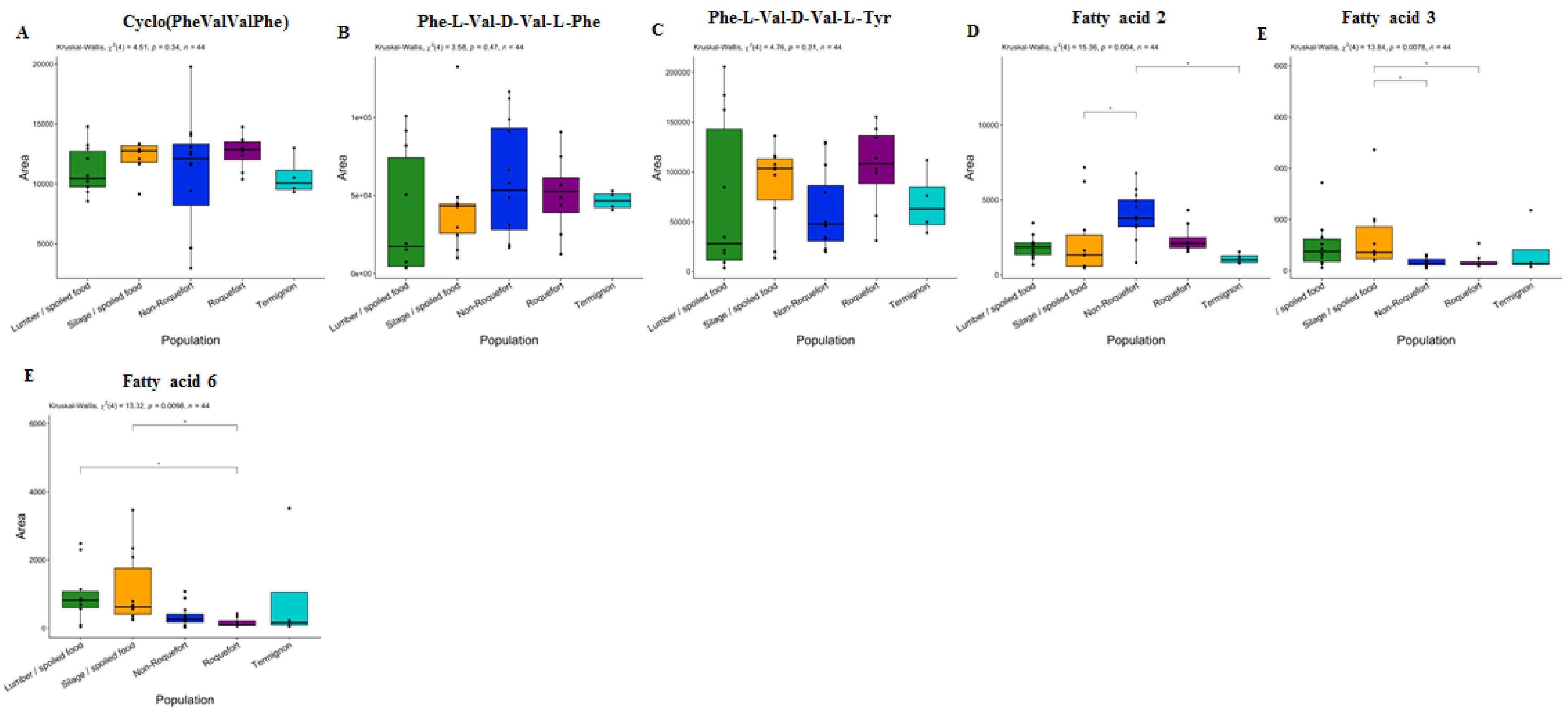


**Supplementary Figure 2: Production level of cyclo-(Phe-Val-Val-Phe) (A), Phe-Val-Val-Phe (B), Phe-Val-Val-Tyr (C), fatty acid 2 (D), fatty acid 3 (E), fatty acid 6 (F), among the five *Penicillium roqueforti* populations.** Production level is expressed as the surface of the peak area of the targeted metabolite per extract matrix mass and mycelium mass. The different populations were colour-coded as follows: green for the lumber/spoiled food population, orange for the silage/spoiled food population, dark blue for the non-Roquefort cheese population, purple for the Roquefort cheese population and light blue for the Termignon cheese population. The results of the global test for a population effect is given at the top of each panel. Pairwise significant differences are indicated by asterisks. The boxplots represent the median (center line), the first quartile and third quartile (box bounds), the maximum and minimum excluding outlier points (whiskers), points being the outliers, *i.e.* with values either below the first quartile minus 1.5 fold the interquartile range or above the third quartile plus 1.5 fold the interquartile range.

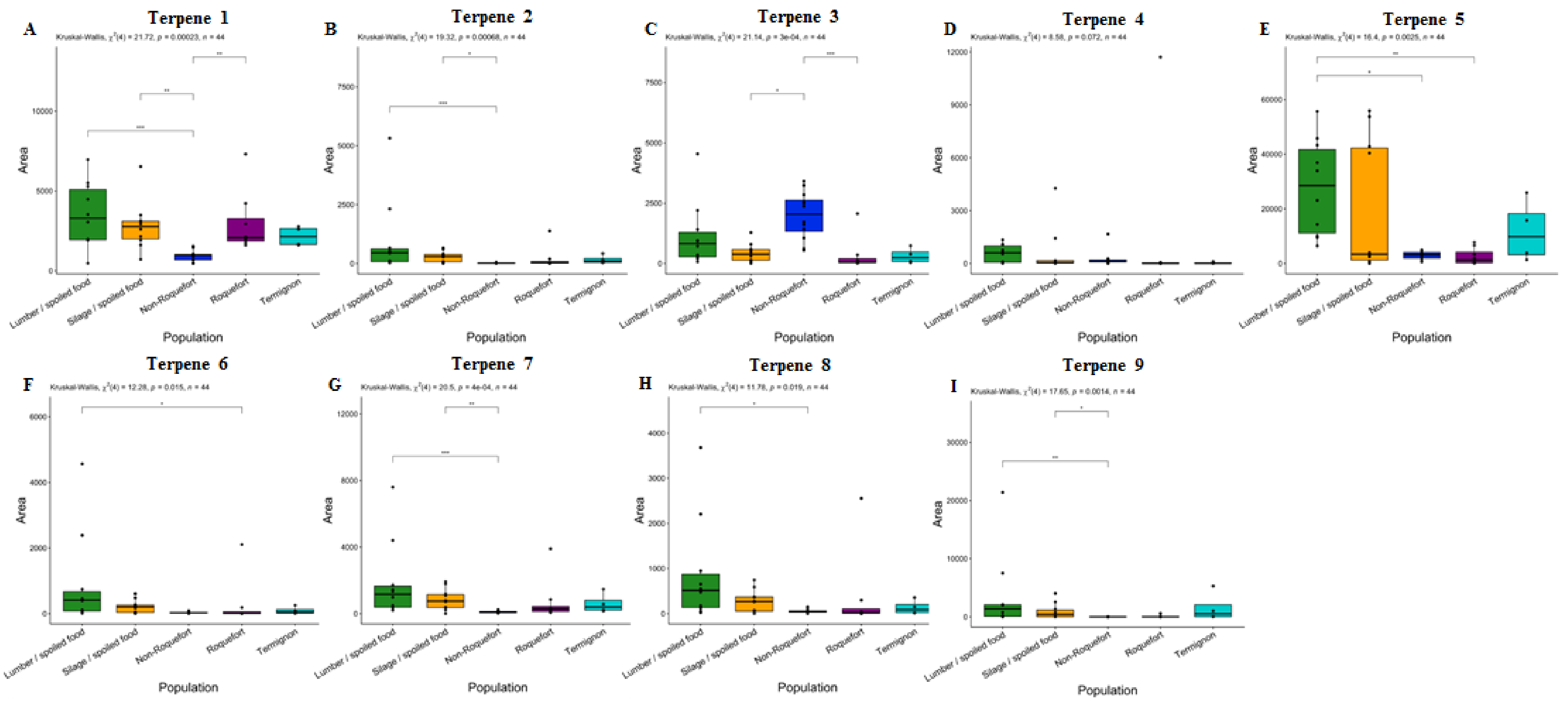


**Supplementary Figure 3: Production level of terpene 1 (A), terpene 2 (B), terpene 3 (C), terpene 4 (D), terpene 5 (E), terpene 6 (F), terpene 7 (G), terpene 8 (H), terpene 9 (I) with formulas in Table 1, among the five *Penicillium roqueforti* populations .** Production level is expressed as the surface of the peak area of the targeted metabolite per extract matrix mass and mycelium mass. The different populations were colour-coded as follows: green for the lumber/spoiled food population, orange for the silage/spoiled food population, dark blue for the non-Roquefort cheese population, purple for the Roquefort cheese population and light blue for the Termignon cheese population. The results of the global test for a population effect is given at the top of each panel. Pairwise significant differences are indicated by asterisks. The boxplots represent the median (center line), the first quartile and third quartile (box bounds), the maximum and minimum excluding outlier points (whiskers), points being the outliers, *i.e.* with values either below the first quartile minus 1.5 fold the interquartile range or above the third quartile plus 1.5 fold the interquartile range.

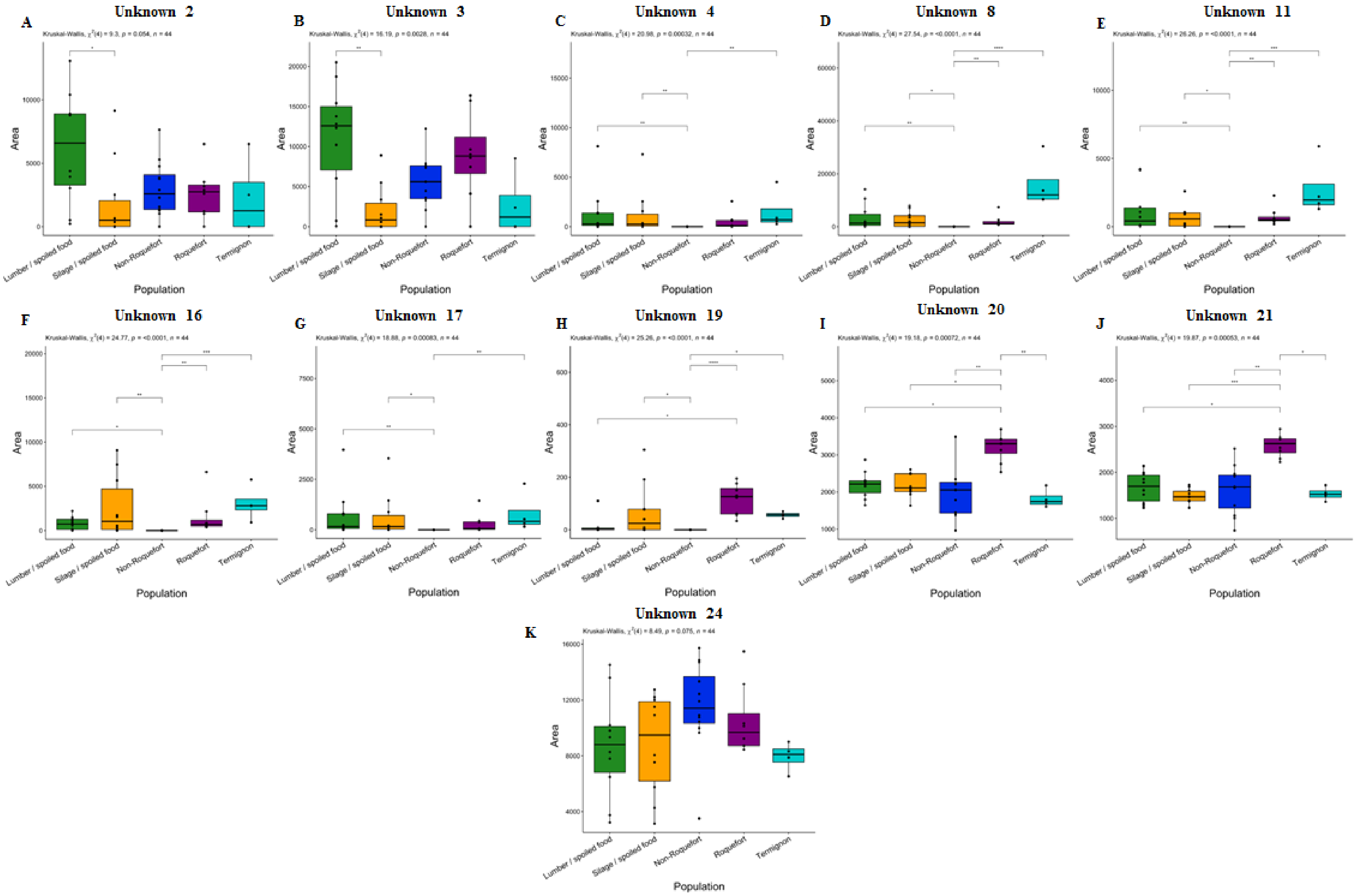

A
Unknown 2
Kruskal-Wallis, χ2(4) = 9.3, p = 0.054, n = 44
B
Unknown 3
Kruskal-Wallis, χ2(4) = 16.19, p = 0.0028, n = 44
C
Unknown 4
Kruskal-Wallis, χ2(4) = 20.98, p = 0.00032, n = 44
D
Unknown 8
Kruskal-Wallis, χ2(4) = 27.54, p = <0.0001, n = 44
E
Unknown 11
Kruskal-Wallis, χ2(4) = 26.26, p = <0.0001, n = 44
F
Unknown 16
Kruskal-Wallis, χ2(4) = 24.77, p = <0.0001, n = 44
G
Unknown 17
Kruskal-Wallis, χ2(4) = 18.88, p = 0.00083, n = 44
H
Unknown 19
Kruskal-Wallis, χ2(4) = 25.26, p = <0.0001, n = 44
I
Unknown 20
Kruskal-Wallis, χ2(4) = 19.18, p = 0.00072, n = 44
J
Unknown 21
Kruskal-Wallis, χ2(4) = 19.87, p = 0.00053, n = 44
K
Unknown 24
Area
Lumber / spoiled food
Silage / spoiled food
Non-Roquefort
Roquefort
Termignon
Population

**Supplementary Figure 4: Production level of unknow 2 (A), unknow 3 (B), unknow 4 (C), unknow 8 (D), unknow 11 (E), unknow 16 (F) unknow 17 (G), unknow 19 (H), unknow 20 (I), unknow  21 (J), unknow 24 (K) with formulas in Table 1, among the five *Penicillium roqueforti* populations.** Production level is expressed as the surface of the peak area of the targeted metabolite per extract matrix mass and mycelium mass. The different populations were colour-coded as follows: green for the lumber/spoiled food population, orange for the silage/spoiled food population, dark blue for the non-Roquefort cheese population, purple for the Roquefort cheese population and light blue for the Termignon cheese population. The results of the global test for a population effect is given at the top of each panel. Pairwise significant differences are indicated by asterisks. The boxplots represent the median (centre line), the first quartile and third quartile (box bounds), the maximum and minimum excluding outlier points (whiskers), points being the outliers, *i.e.* with values either below the first quartile minus 1.5 fold the interquartile range or above the third quartile plus 1.5 fold the interquartile range.
